# Primate neuronal connections are sparse as compared to mouse

**DOI:** 10.1101/2020.09.24.311852

**Authors:** G.A. Wildenberg, M.R. Rosen, J. Lundell, D. Paukner, D.J. Freedman, N. Kasthuri

**Affiliations:** Department of Neurobiology, University of Chicago, Chicago, IL, USA; Argonne National Laboratory, Lemont, IL, USA

## Abstract

The mouse and macaque primary visual cortices are foundational models of cortical functioning, particularly at the level of single neurons. Therefore, detailing differences in how individual neurons connect across these species would inform models of cortical functioning and of how brains evolve. However, existing comparisons are limited, measuring synapse density without regard to where synapses are made or on what types of neurons. We use large volume electron microscopy to address this gap, reconstructing a total of 7735 synapses across 160 total neurons (146 excitatory, 14 inhibitory) from adult Rhesus macaque and mouse Layer 2/3 of primary visual cortex (V1). We find that primate connections are broadly sparse: primate excitatory and inhibitory neurons received 3-5 times fewer spine and somatic synapses with lower ratios of excitatory to inhibitory synapses than mouse equivalents. However, despite reductions in absolute synapse number, patterns of axonal innervation were preserved: inhibitory axons sparsely innervated neighboring excitatory neurons in macaque and mouse at similar rates and proportions. On the output side, most excitatory axons in mice myelinated close to the soma (81%) while most primate axons (68%) did not. Interestingly, primate axons, but not mouse axons, that myelinated had 3.3 fold more axon initial segment synapses than axons that did not myelinate, suggesting differential inhibitory control of long distance output in primate brains. Finally, we discover that when artificial recurrent neural networks (RNNs) are constrained by the metabolic cost of creating and maintaining synapses, increasing the number of nodes (e.g. neurons) as networks optimize for a cognitive task, reduces the number of connections per node, similar to primate neurons as compared to mice.

**One Sentence Summary:** Using large volume serial electron microscopy, we show that primate cortical neural networks are sparser than mouse and using recursive neural nets, we show that energetic costs of synaptic maintenance could underlie this difference.

## Introduction

The two dominant models in neuroscience for understanding cortical processing at the level of individual neurons are the primary visual systems of *mus musculus* and *macaca mulatta*, with expansive literatures on morphological, functional, and molecular properties of neuronal classes (Bakken et al., 2016; Bernard et al., 2012; Gilman et al., 2017; Gouwens et al., 2018; Gouwens et al., 2019; Gur and Snodderly, 2008; Medalla and Luebke, 2015; Ohki et al., 2005; Tasic et al., 2016; Tasic et al., 2018). The advantages of each are obvious: the small size and genetic manipulability of the mouse nervous system and the close homology of macaque brains and behavior to humans and comparisons across both would leverage these advantages to better understand how neuronal cell types change across brains of such disparate size and neuronal number (Herculano-Houzel, 2009, 2011). However, while molecular and functional comparisons continue apace (Bakken et al., 2020; Hodge et al., 2019; Krienen et al., 2020; Wang et al., 2016), there is little known about how classes of neurons in mouse and primate brains physically connect with each other: are there species differences in the distributions of excitatory and inhibitory inputs on neurons of different types? How are long and short distance outputs of individual neurons organized? Do basic patterns of neuronal connections differ? A better understanding of these differences would suggest potential mechanisms of how connections on individual neurons change as brains scale in size and neuronal number.

Here we detail how excitatory and inhibitory neurons in adult macaque and mouse primary visual cortices differ in the numbers and types of connections received and how those differences affect basic properties of connectivity (e.g. the frequency with which neighboring neurons share inhibitory inputs). Finally, we use neurobiologically inspired Artificial Neural Networks (ANNs) constrained with metabolic costs to explore potential mechanisms for the changes in neuronal connections we see across species.

We applied recent advances in large volume, automated serial electron microscopy and computational analyses-*e.g.* ‘connectomics’ (Baena et al., 2019; Briggman et al., 2011; Gour et al., 2020; Hayworth et al., 2020; Januszewski et al., 2018; Turner et al., 2020b; Vishwanathan et al., 2020; Yin et al., 2019) to compare the numbers and types of connections received by individual identified neurons in the primary visual cortex (V1) of mouse (*Mus musculus*) and primate (*Macaca mulatta*). We used a connectomic approach since many techniques for mapping long and short-range connections in the mouse (Bae et al., 2018; Karimi et al., 2020; Morgan and Lichtman, 2020; Motta et al., 2019; Oh et al., 2014) are not widely available in the primate and electron microscopy remains the gold standard for identifying neuronal connections. While there is a long history of pioneering studies using electron microscopy in this regard (DeFelipe et al., 2002; Gilman et al., 2017; Hsu et al., 2017; Sherwood et al., 2020), until recently, electron microscopic reconstructions at the scale of individual neurons remained technically and computationally intractable (Baena et al., 2019; Hayworth et al., 2020; Januszewski et al., 2018; Kasthuri et al., 2015; Motta et al., 2019; Yin et al., 2019). Thus, existing studies primarily focus on differences in the number or size of synapses across species, *largely without knowing the type of neuron making or receiving these connections.* For example, it remains unknown how synaptic density changes reflect changes in excitatory and inhibitory synapses on individual inhibitory or excitatory neurons and/or on different parts of neurons (e.g. on dendrites vs. soma vs. axon initial segment). Finally, data about inhibitory connections and connections made onto inhibitory neurons remain scarce, since such data cannot be easily inferred from light level reconstructions of neurons (i.e. Golgi staining and spine counting).

We find that primate neural networks are sparse relative to mice: primate excitatory and inhibitory neurons receive 3 to 5-fold fewer excitatory and inhibitory inputs but the patterns of connections of inhibitory axons are preserved. Using neurobiologically-inspired artificial neural networks constrained by metabolic costs and trained on cognitive tasks, we find that as neural networks grow (i.e. add more nodes or ‘neurons’), the energy constraints of building and maintaining synapses produces larger networks with reduced numbers of connections per node.

## Results

We prepared 300µm thick sections of ‘adult’ primary visual cortex (V1) from mouse (15 weeks old, male) and from the foveal region (see Methods) of two primates (11 and 14.5 years old, male) for multi-scale EM-based connectomics. Because comparing ages across species is difficult, we chose ages past a majority of brain developmental milestones but before onset of age-associated pathologies for both species (Bakken et al., 2016; Bourgeois and Rakic, 1993; Horton and Hocking, 1997; Lee et al., 2000; Peters et al., 1996; Scott et al., 2016; Semple et al., 2013). V1 was identified in each species using areal and cellular landmarks (Fig. S1, A and B), and in primates, the foveal region was selected (Van Essen et al., 1984). Samples were prepared as previously described (Hua et al., 2015) (and see **Methods**). We first collected 3,000 ultra-thin sections using the ATUM approach (Kasthuri et al., 2015) from mouse (each section: 0.8mm x 1.5 mm x 40nm) and 3,000 from primate (0.8mm x 2.4mm x 40nm), where each individual section spanned all cortical layers in both species (**Fig. 1, A and B**). We imaged the depth of cortex with low resolution EM (∼40 nm in plane resolution) and skeletonized the dendritic arbors of 102 primate neurons and 68 mouse neurons across all cortical layers (**Fig. 1, A and B**). We used these reconstructions to evaluate neuronal shapes and densities between species and identify layers (L2/3) for synaptic level reconstructions (movies S1 and S2). We restricted our subsequent cellular and synaptic analyses to Layer 2/3 since, 1) neurons in both species have smaller dendritic arbors (Rojo et al., 2016) (e.g. as compare to dendritic arbors Layer 5 neurons), allowing us to capture a larger fraction of all the synapses on a particular neuron, and 2) there seems to be a limited number of morphological cell types in L2/3 (Gouwens et al., 2019; Radnikow and Feldmeyer, 2018; Wang et al., 2018) making for potentially cleaner comparisons across species.

**Fig. 1.**
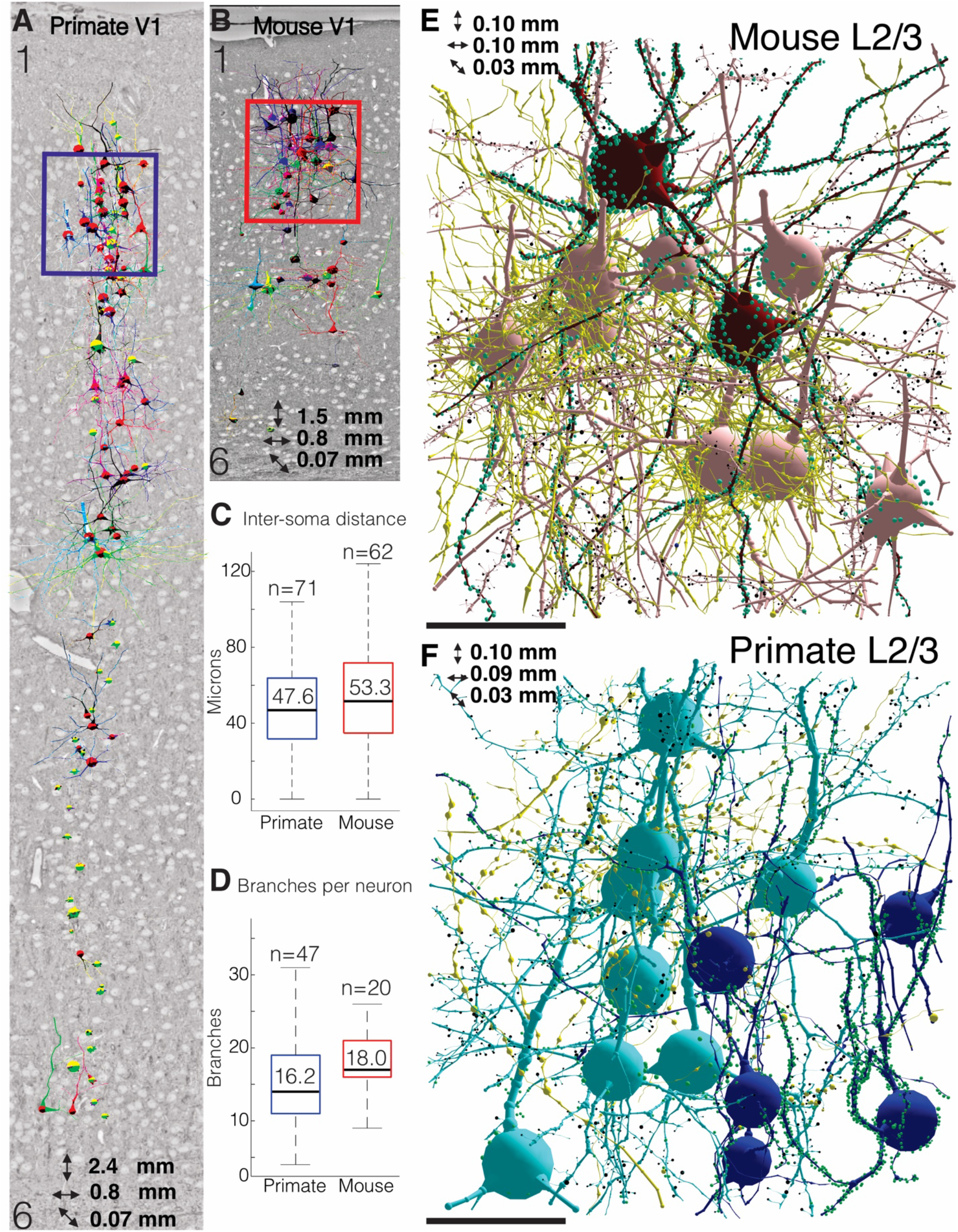
Similar meso-scale architectures of primary visual cortex (V1) in macaque and mouse. **(A-B)** Reconstructions of 102 primate (A) and 68 mouse (B) neurons from primary visual cortex (V1), superimposed in place on single EM sections spanning layers 1-6. Blue and red boxes indicate regions of fine resolution imaging. (**C**) Box-and-whisker plot of distances between L2/3 neuronal soma in primate and mouse (primate: 47.6 ± 0.29 µm, n = 71; mouse: 53.3 ± 0.38 µm, n = 62, P=2.8e-3). (**D**). Box-and-whisker plot of dendritic branch number of L2/3 neurons (primate: 16.2 ± 1.2, n= 47; mouse: 18.0 ± 0.9, n= 20, P=0.08). (**E-F**). 3D reconstructions of representative neurons from mouse (0.10 x 0.10 x 0.03 mm) and primate (0.10 x 0.09 x 0.03 mm) V1, L2/3 sub-volumes re-imaged at fine (∼ 6nm XY) resolution. Excitatory neurons (+Neurons) are light red or blue and inhibitory neurons (-Neurons) are dark red or blue in mouse and primate, respectively. Green nodes are inhibitory synapses on shafts and soma, black nodes are excitatory synapses on dendritic spines, fine yellow skeletons are axons. Scale bar = 50 µm. Data: mean ± SEM. *P*-values: two-tailed Mann-Whitney U test.

We first looked for differences in the cytoarchitecture of L2/3 between species since an advantage of large volume EM is that all cells are completely labeled in the volume, unlike sparse labeling approaches like Golgi staining (Vints et al., 2019). We found that excitatory somata were distributed at similar densities across both species (**Fig. 1C**; P = 2.8e-3, Mann-Whitney), had similar somatic surface areas (**Fig. S2A**; P=0.3, Mann-Whitney), and number of dendritic branch points (**Fig. 1D**; P = 0.08, Mann-Whitney). Finally, we found that the total length of dendrite per neuron was larger in mouse (**Fig. S2B**; P = 1.0e-5, Mann-Whitney), similar to previous reports (Gilman et al., 2017). Visual inspection of the neurons (**Fig. S3**) revealed that the majority of neurons in both species were pyramidal with apical pointing dendrites, and that 3/37 soma in the mouse and 9/42 primate soma were inhibitory neurons, agreeing with previously reported ratios (Džaja et al., 2014; Sultan and Shi, 2018).

Given their similarity of shape and distributions, we next analyzed whether mouse and primate neurons differed in the number and types of neuronal connections received. We annotated 3,973 synapses and 186 axons that synapsed onto 64 excitatory and 4 inhibitory neurons in the mouse and 3,762 synapses and 94 axons that synapsed on 82 excitatory and 10 inhibitory neurons across two primates. **Figure 1, E and F** show representative examples of mouse and primate neurons we reconstructed. We analyzed the connectivity of different neuronal compartments (e.g. dendrites, soma, and axon initial segments) across species. We classified each synapse as a spine, shaft, somatic, or perisomatic (Freund and Katona, 2007), and each axon as excitatory or inhibitory based on whether they predominantly innervated spines, or shafts and somas, respectively (Harris and Weinberg, 2012). For dendrites, we counted spine and shaft synapses at different dendritic locations relative to the soma, in order to account for synapse variability within a neuron’s dendritic tree (Ballesteros-Yáñez et al., 2006; Benavides-Piccione et al., 2013; Vaudin et al., 1988).

### Somatic and Dendritic Innervation of Primate and Mouse Neurons

We discovered a broad reduction in the number of both excitatory and inhibitory types received by primate inhibitory and excitatory neurons relative to the mouse. Figure 2 depicts four representative examples of excitatory and inhibitory primate and mouse neurons and all the connections they received while Figure 3 shows statistical quantification across multiple neurons and across multiple samples which we also summarize in **Table S1**.

**Fig. 2.**
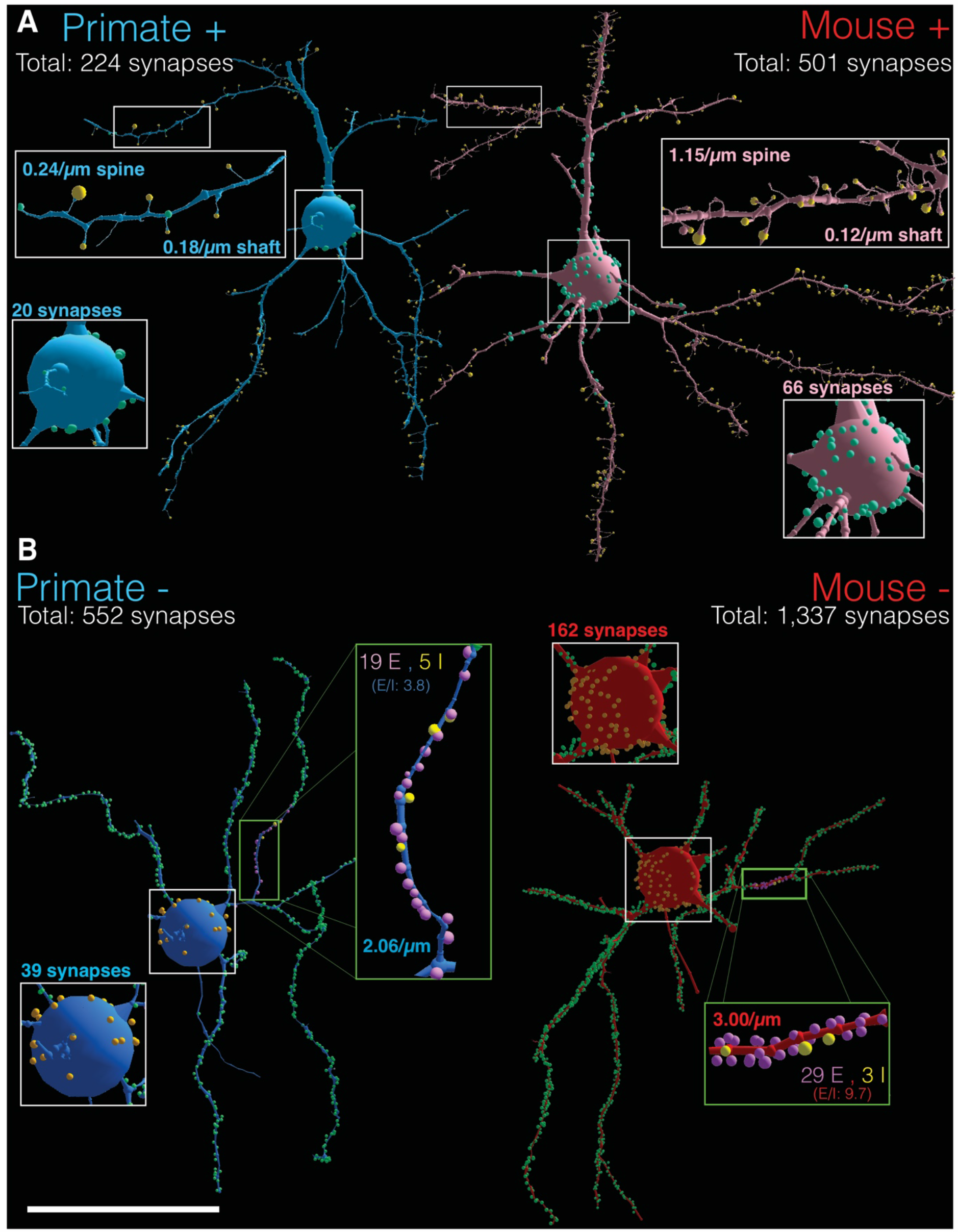
Mouse L2/3 excitatory and inhibitory neurons receive more connections than primate neurons. (**A**) Representative EM reconstructions of morphologically matched L2/3 primate and mouse excitatory (+) neurons. All synapses were annotated and quantified (soma: total # of synapses, dendrite: spine synapses/µm; shaft synapses/µm). *Square inset*: magnified views of soma: somatic synapses are green; Primate + neuron: 20 somatic synapses; Mouse + neuron: 66 somatic synapses. *Rectangle Inset*: magnified views of dendrites: shaft synapses are green spheres and spine synapses are yellow spheres; Primate + neuron: 0.24 spine synapses/µm, 0.18 shaft synapses/µm; Mouse + neuron: 1.15 spine synapses/µm, 0.12 shaft synapses/µm. (**B**) Representative EM reconstructions of morphologically matched L2/3 primate and mouse inhibitory (-) neurons. All synapses were annotated and quantified (soma: total # of synapses, dendrite: spine synapses/µm). *Square inset*: magnified views of soma: somatic synapses are orange; Primate - neuron: 39 somatic synapses; Mouse - neuron: 162 somatic synapses. *Rectangle Inset*: magnified views of dendrites: excitatory shaft synapses are pink spheres and inhibitory shaft synapses are yellow spheres; Primate – neurons: 2.06/µm shaft synapses/µm; 19 excitatory synapses and 5 inhibitory: E/I = 2.8; Mouse – neurons: 3.00 shaft synapses/µm; 29 excitatory synapses and 3 inhibitory: E/I = 9.7. Scale Bar = 45 µm.

**Fig. 3.**
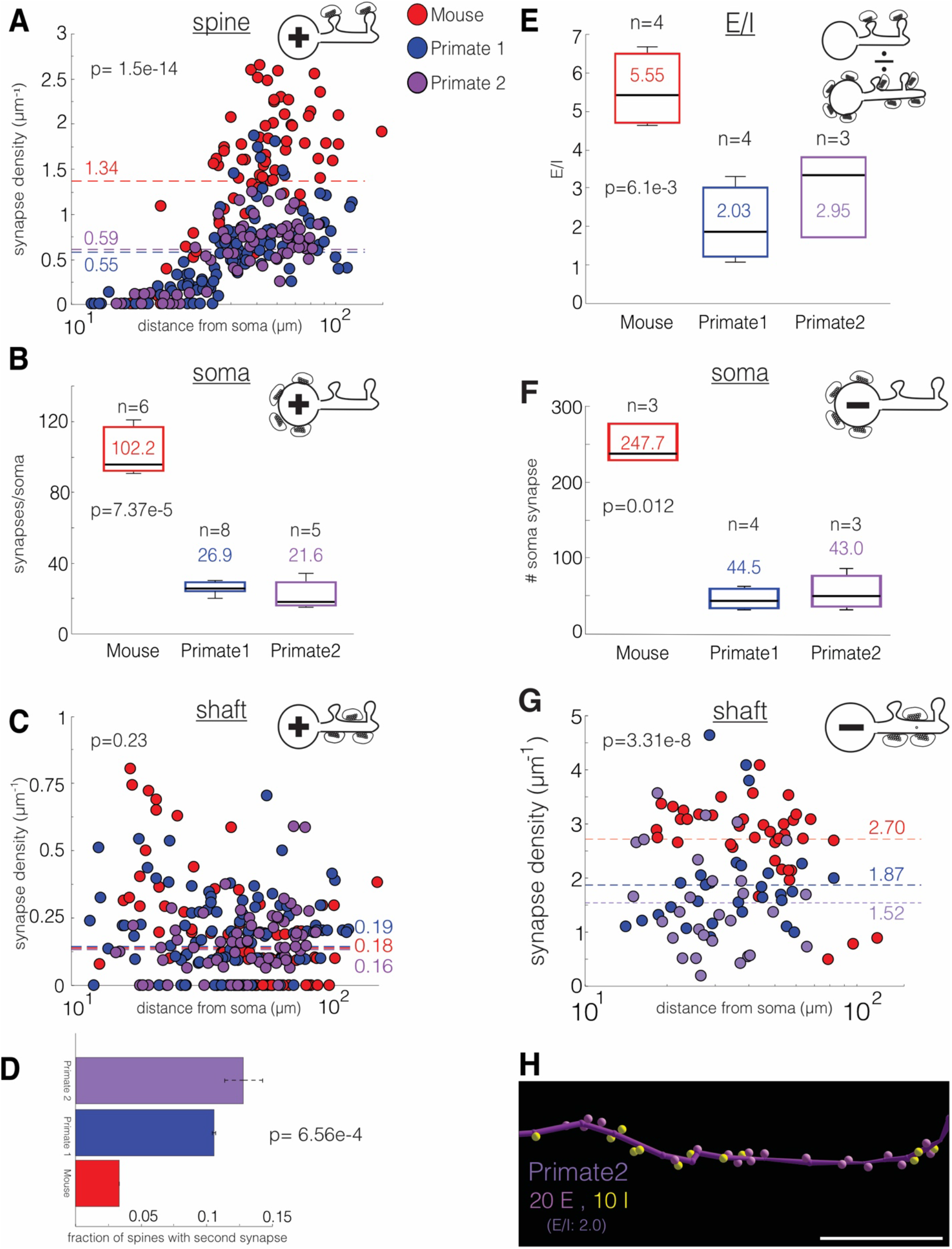
Increased connectivity in mouse L2/3 excitatory and inhibitory neurons is common. All data is from mouse (red circles), and two primate (blue and purple circles) L2/3 datasets. (**A**) Scatter plot of excitatory neuron spine synapses/µm versus distance from soma (µm) (primate 1: 0.55 ± 0.04 synapses/µm, n = 109 10 µm dendrite fragments across 10 neurons; primate 2: 0.59 ± 0.04 synapses/µm, n = 66 10 µm dendrite fragments across 6 neurons; mouse: 1.34 ± 0.09 synapses/µm, n = 82 10µm dendrite fragments across 10 neurons; P=1.5e-14). (**B**) Box-and-whisker plot of excitatory neuron synapses/soma (mouse: 102.2 ± 5.3, n = 388 synapse across 6 soma54; primate 1: 26.9 ± 2.0, n = 129 synapses across 8 soma; primate 2: 21.6 ± 3.44, n = 108 synapses across 5 soma; P=7.37e-5). (**C**) Scatter plot of excitatory neuron shaft synapses/μm versus distance from soma (μm) in excitatory neurons (primate 1: 0.19 ± 0.02 synapses/μm, n = 109 10 μm dendrite fragments counted across 10 neurons; primate 2: 0.16 ± 0.01 synapses/µm, n = 66 10 µm dendrite fragments across 6 neurons; mouse: 0.18 ± 0.02 synapses/μm, n = 82 10 μm dendrite fragments counted across 10 neurons; P=0.23). (**D**) Primate excitatory neurons have more 2^nd^ synapses. bar plot of the fraction of spines with 2^nd^ synapses (primate 1: 0.11 ± 0.001, n = 180 spines across 3 neurons; primate 2: 0.13 ± 0.01, n = 257 spines across 4 neurons; mouse: 0.03 ± 0.00, n = 180 spines across 3 neurons; P = 6.56e-4). **(E)** Box-and-whisker plot of the excitatory to inhibitory synapse (E/I) ratio on excitatory neurons (see Methods) (mouse: 5.55 ± 0.48, n= 770 spine synapses (180 scored for 2^nd^ spine synapses), 109 shaft synapses, and 401 somatic/perisomatic synapses over 4 neurons; primate 1: 2.03 ± 0.48, n = 427 spine synapses (180 scored for 2^nd^ spine synapses), 151 shaft synapses,106 somatic/perisomatic synapses over 4 neurons; primate 2: 2.95 ± 0.63, n = 242 spine synapses (197 scored for 2^nd^ spine synapses), 56 shaft synapses, 67 somatic/perisomatic synapses over 3 neurons; P = 6.1e-3). **(F)** Box-and-whisker plot of inhibitory neuron synapses/soma (mouse: 247.7 ± 14.8, n = 743 synapses across 3 soma; primate 1: 44.5 ± 6.7, n = 178 synapses across 4 soma; primate 2: 43.0 ± 14.6, n = 129 synapses across 3 soma; P=0.012). **(G)** Scatter plot of inhibitory neuron shaft synapses/µm versus distance from soma (µm) (primate 1: 1.87 ± 0.17 synapses/µm, n = 29 10µm dendritic fragments across 5 neurons; primate 2: 1.52 ± 0.17 synapses/µm, n = 29 10µm dendritic fragments across 5 neurons; mouse: 2.70 ± 0.12 synapse/µm, n = 39 10µm dendrite fragments across 4 neurons; P=3.31e-8). **(H)** Reconstruction of excitatory (purple spheres) and inhibitory (yellow spheres) synapses along the dendrite of an inhibitory neuron from the second primate. The second primate has 20 excitatory synapses for every 10 inhibitory (E/I = 2.0). Scale bar = 10 µm Data: mean ± SEM. *P*- values: two-tailed Mann-Whitney U test.

The primate excitatory neuron had ∼three-fold lower dendritic spine synapses relative to the mouse and throughout the dendritic arbor (**Fig 2A**, dendrite insets, primate, 0.24 excitatory spine synapses/µm; mouse, 1.15 excitatory spine synapses/µm). Similarly, somatic inputs were reduced three-fold: the example primate soma received 20 somatic and perisomatic synapses (henceforth collectively called “somatic synapses”) while the example mouse soma received 66 somatic synapses (**Fig 2A**, soma inset). We found two classes of synapses where innervation on primate neurons was the same or more prevalent than mouse: 1. shaft synapses (*i.e.* synapses on dendritic shafts between spines) were less prevalent than spine synapses in both species, but primate and mouse neurons had similar densities, unlike spine or somatic synapses (0.18 shaft synapses/µm for primate, 0.12 shaft synapses/µm for mouse). 2. While primate excitatory neurons receive three-fold *fewer* spine synapses, individual spines of primate neurons were three-fold *more* likely to receive a second, inhibitory synapse (**Fig. S4A-B**) (Jones and Powell, 1969; Knott et al., 2002; Kwon et al., 2019).

We saw a similar trend in inhibitory neurons. In the same volume (Fig. S3), we identified inhibitory neurons based on the relative sparsity of spines on their dendrites ((Freund and Buzsáki, 1996; Gulyás et al., 1999) and see results) and found that primate inhibitory neurons received ∼1.5x fold fewer shaft synapses than mouse inhibitory neurons and 4-fold fewer somatic synapses (**Fig. 2B**, dendrite and somatic insets: primate inhibitory neuron - 2.06 shaft synapses/µm and 39 somatic synapses compared to 3.00 shaft synapses/µm and 162 somatic synapses in the mouse). We classified shaft synapses on inhibitory dendrites as excitatory or inhibitory by tracing 24 axons in the primate, and 32 axons in the mouse that made shaft synapses on similar regions of dendrites of matched inhibitory neurons (**Fig. 2B**, green boxes) and characterized every additional synapse those axons made in the volume as excitatory (E, purple spheres-predominantly spine synapses) or inhibitory (I, yellow spheres-predominantly shaft and soma synapses; see Methods). We thus calculated for matched inhibitory dendritic segments that the mouse inhibitory neurons received 29 excitatory inputs and 3 inhibitory inputs and the primate inhibitory neuron received 19 excitatory inputs and 5 inhibitory inputs (**Fig. 2B**). Like excitatory neurons, we find primate inhibitory neurons received fewer excitatory and somatic connections but similar numbers of inhibitory dendrite synapses relative to mouse.

We found statistically significant results across multiple excitatory and inhibitory neurons over multiple samples (n=26 excitatory and 14 inhibitory neurons across 1 mouse and 2 primates, (see Table S1 for summary). Primate neurons received ∼ 3-fold fewer spine synapses (Fig 3A; P = 1.5e-14, Mann Whitney) and 4-fold fewer somatic synapses (Fig 3B, P=7.37e-5, Mann Whitney) than similar mouse excitatory neurons. The numbers of shaft synapses where comparable (**Fig. 3C**; P = 0.23, Mann Whitney) and individual primate spines where 3-fold more likely to be innervated by a second inhibitory axon (**Fig. 3D**; P = 6.56e-4, Mann-Whitney). Indeed, while individual excitatory neurons within a particular species varied in the number of connections they receive, such variability was much smaller than the differences we found across species (**Fig. S5).** Using these spine, shaft, somatic, and second spine averages and mean dendritic arbor lengths for primate and mouse layer 2/3 neurons as previously reported (Gilman et al., 2017), we calculated the ratio of the total number of excitatory and inhibitory inputs onto individual excitatory neurons (see **Methods**). We estimated that mouse neurons received on average 6894 total connections (5847 excitatory and 1048 inhibitory) and across the two primate samples, we estimated that primate excitatory neurons received on average 2499/2617 total connections (1698/1914 excitatory and 802/703 inhibitory; Primate1/Primate2). Thus, primate excitatory neurons had lower E/I ratios than mouse neurons (**Fig. 3E**; P = 6.1e-3, Man-Whitney).

We see similar reductions across multiple inhibitory neurons. Primate inhibitory neurons receive less somatic synapses (**Fig. 3F**; P=0.012, Mann-Whitney), and reduced shaft synapse density (**Fig. 3G**; P=3.31e-8, Mann-Whitney) relative to mouse inhibitory neurons. Among dendritic synapses on primate inhibitory neurons, we observed a similar increase in the proportion of inhibitory inputs onto inhibitory neurons in the second primate dataset (**Fig 3H**; 20 excitatory inputs for every 10 inhibitory inputs; E/I = 2.0). Thus, we find a dramatic global and statistically significant reduction in the numbers of excitatory and inhibitory inputs received by both primate excitatory and inhibitory neurons.

### Axonal Myelination and Innervation of Primate and Mouse Neurons

We next examined whether synapses onto the axons of primate excitatory neurons showed similar reductions in synapse number (summarized in Table S2). We first classified L2/3 excitatory neurons in both species into those with a non-branching, basally oriented axon that myelinated within ∼25 µm from the soma (MA), and a second class whose axons formed local collaterals and left the field of view unmyelinated (uMA; **Figure. 4A**). For every axon, we measured the axonal initial segment (AIS) (i.e. the distance from the soma to the most distal axo-axonic synapse) and the total number of axo-axonic synapses along that length. We found significant species differences: 1. mouse L2/3 excitatory neurons were far more likely to myelinate (13/16) in our volumes than primate neurons (11/23) (**Fig. 4A, bottom inset** P = 0.003, Mann-Whitney) and 2. while there was little difference in the length of the AIS across species and cell type (**Fig. 4C**; P = 0.02, Mann-Whitney), primate axons that myelinated received ∼4 fold more AIS synapses than primate axons that did not myelinate (**Fig. 4D**; P = 1.54e-7, Mann-Whitney) while AIS innervation of mouse neurons was independent of their myelination status (**Fig 4E**; P=0.08, Mann-Whitney). Thus, we found two differences between primate excitatory axons and mouse: a higher rate of myelination correlated with a significant increase in synapses along axon initial segments of MA but not uMA primate axons. This correlation increased our confidence that the differences seen in myelination rates between primate and mouse (Fig. 4A, bottom panel) are biologically relevant and unlikely due to primate neurons axons leaving the imaged volume before we detected a myelination event.

**Figure 4.**
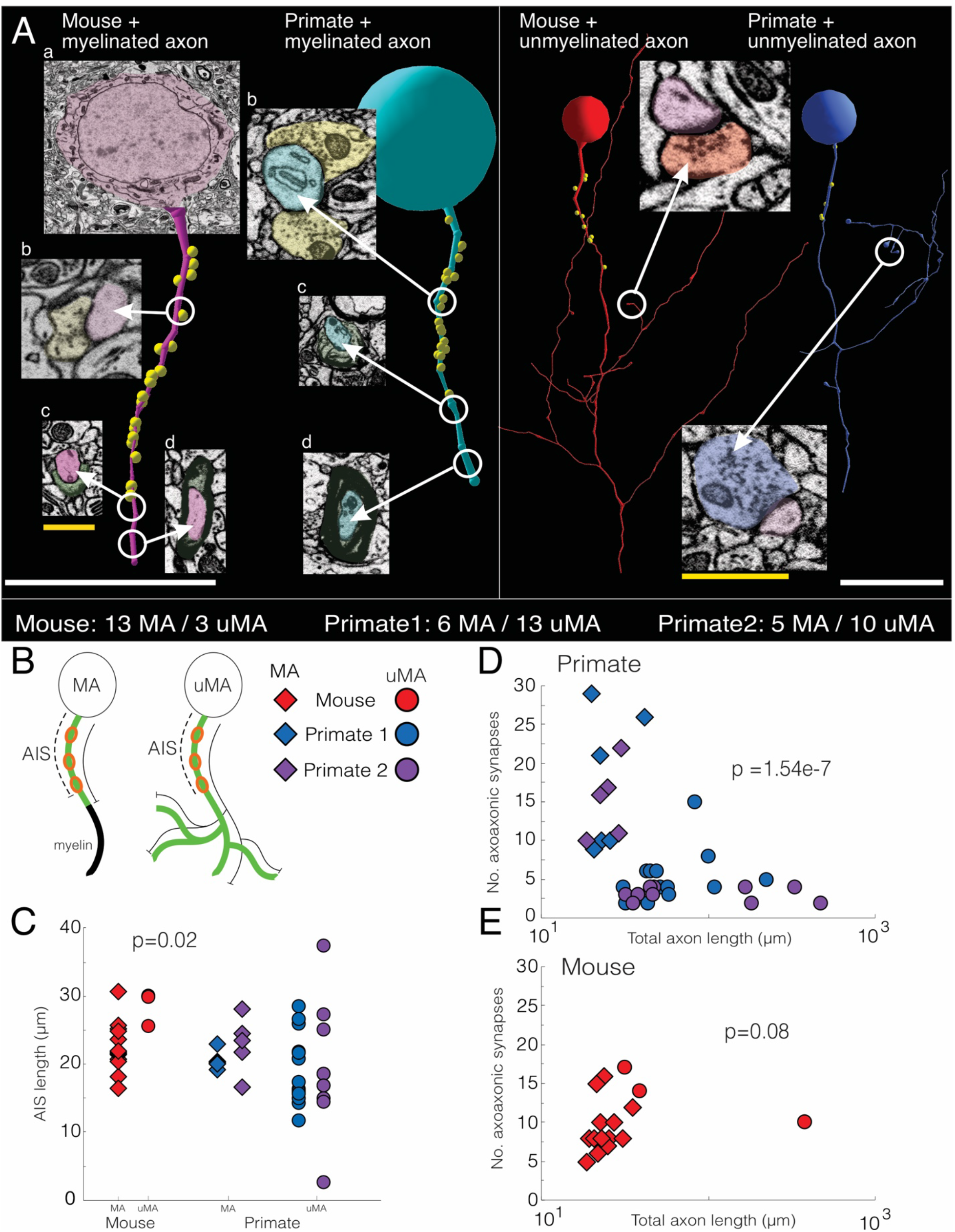
Mouse and primate L2/3 excitatory neurons differ in their axon profiles. **(A)** Representative reconstructions of Mouse and Primate L2/3 myelinated and unmyelinated excitatory neurons. Left panel: mouse (pink) and primate (light blue) L2/3 excitatory (+) neurons possessing a non-branching, basally directed axon that myelinates ∼25 µm from the soma and leaves the field of view as a myelinated axon. Yellow nodes in reconstructions represent locations of axo-axonic synapses. 2D EM images depict: (a) soma, (b) points of axo-axonic synapses (yellow synapses onto red or light blue axon), (c) point of initial myelination, and (d) fully myelinated axon. Right panel: mouse (red) and primate (dark blue) L2/3 excitatory (+) neurons with unmyelinated axons that form local collaterals and synapses. Yellow nodes in reconstructions represent locations of axo-axonic synapses. 2D EM images depict synapses made by mouse and primate axon collaterals (red and blue for mouse and primate, respectively) onto a post-synaptic target (purple). *Bottom panel*: ratio of excitatory neurons with myelinated (MA) to unmyelinated (uMA) in mouse and both primate datasets. Ratio between mouse and primate is statistically significant: P=0.003. **(B)** Cartoon describing our quantification schemes for excitatory neuron myelinated (MA) and unmyelinated axons (uMA): (1) the distance between the soma and furthest axoaxonic synapses was reported as the axonal initial segment (AIS) (dotted black line) and (2) the total number of axoaxonic synapses (open red circles) per total axon length (solid black line). Data is depicted in C-E as follows: MA (diamond) and uMA (circle) populations for mouse (red) and primate samples (blue, purple). **(C)** Scatter plot of the AIS for mouse and primate uMA and MA excitatory neurons (mouse uMA: 28.4 ± 1.5 µm, n = 3 neurons; mouse MA: 22.5 ± 1.0 µm, n = 13 neurons; primate 1 uMA: 19.3 ± 1.5 µm, n = 13 neurons; primate 1 MA: 20.4 ± 0.5 µm, n= 6 neurons; primate 2 uMA: 19.2 ± 3.0 µm, n= 10 neurons; primate 2 MA: 22.8 ± 2.0 µm, n= 5 neurons; P = 0.02). (**D-E)** Scatter plots of lengths of un-myelinated (uMA) and myelinated (MA) axons versus the number of axoaxonic synapses the axon receives in primate and mouse. (D) (primate_1 uMA: 5.3 ± 1.0 axoaxonic synapses, n = 69 synapses across 13 neurons; primate_1 MA: 17.5 ± 3.7 axoaxonic synapses, n = 105 synapses across 6 neurons; P1 = 4.4e-4. primate_2 uMA: 3.1 ± 0.3 axoaxonic synapses, n = 31 synapses across 10 neurons; primate_2 MA: 15.2 ± 2.1 axoaxonic synapses, n = 76 synapses across 5 neurons; P2 = 6.7e-4. (E) (mouse uMA: 13.7 ± 2.0 axoaxonic synapses, n = 41 synapses across 3 neurons; mouse MA: 9.3 ± 1.0 axoaxonic synapses, n = 121 synapses across 13 neurons; P = 0.08) and primate. Scale bar: white = 20 µm, yellow = 1 µm. Data: mean ± SEM. *P*-values: two-tailed Mann-Whitney U test.

### Inhibitory Innervation Patterns of Primate and Mouse Neurons

We next focused on neuronal wiring, on how these changes in numbers of connections on individual neurons correlated with how neurons connect with each other. We focused on reconstructing the innervation patterns of axons synapsing with the soma of excitatory neurons, since we could reasonably assume that were inhibitory, likely from parvalbumin + interneurons (Karube et al., 2004; Kisvárday et al., 1985; Somogyi, 1989). Despite the large differences in the absolute numbers of somatic connections, the basic properties of mouse and primate inhibitory axons (i.e. axonal branching, bouton size, and total synapse frequency) were similar between species (**Fig. S6, A-C**). Somatic innervating axons in both species made similar numbers of synapses per soma (**Fig. S6D; mouse: 2.34, primate: 1.33. P = 0.07. Mann-Whitney**), so the ∼4 fold increase in somatic synapse number on mouse excitatory neurons was primarily the result of more axons innervating each somas (**Fig. S6E**; P = 2.1e-3, Mann-Whitney).

We hypothesized that one likely consequence of more axons innervating each mouse soma was that mouse axons were more likely to innervate adjacent soma (i.e. mouse excitatory neurons ‘share’ innervation from individual inhibitory axons). We traced every axon innervating the somas of nearest neighbor excitatory neurons in both species and analyzed the number of inhibitory axons ‘shared’ with the immediately neighboring soma. Despite the large difference in absolute numbers, the fraction of somatic axons that innervated both soma – ‘shared’ - were equivalent, if not slightly larger, in the primate: 7/19 axons shared, 37% in primate and 14/57 axons shared, 25% in mouse (**Figure 5A**). We observed this trend broadly: inhibitory axons were ‘shared’ by neighboring excitatory neurons in primate brains at similar, if not slightly higher, rates than in mouse (**Fig. 5B**; P = 0.1, Mann-Whitney). We extended this analysis beyond nearest-neighbor neurons to ask: how often do axons innervating a particular soma also synapse with the other excitatory neurons in the volume as a function of somatic distance (*cartoon*, **Fig. 5C**). We identified every synapse (spine, shaft, somatic, or perisomatic) these axons made, starting from a central neuron (n= 4 neurons, 2 primates - P1 and P2- and 2 mouse- M1 and M2), with every other excitatory neuron in the volume. We plotted the number of axons shared between central neurons and other neurons versus distance and found a similar decrease in axonal sharing with distance across species (**Fig. 5C**). Neurons from both species showed a sharp decrease in sharing: by 70 µm distance from a central soma, few if any axons were shared with adjacent neurons in either species. Thus, we were unable to disprove our hypothesis that changes in the absolute numbers of connections would correlate with differences in wiring across species. Instead, our results point to a conservation of the basic principles of inhibitory wiring even with a substantially higher number of inhibitory axons innervating each excitatory mouse neuron compared to the primate.

**Fig. 5.**
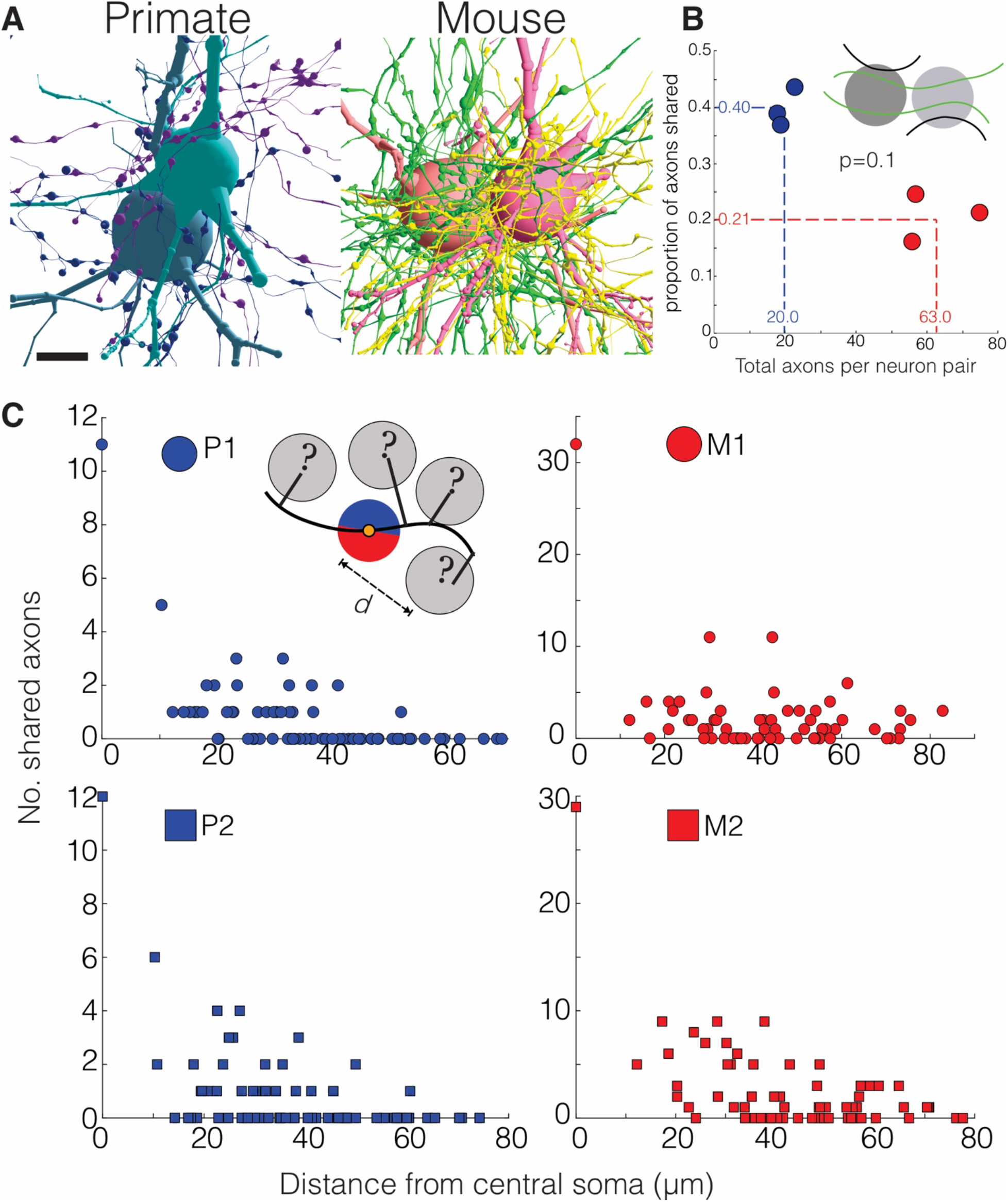
Neighboring primate and mouse excitatory neurons have similar inhibitory networks. (**A**) Reconstructions of every inhibitory axon innervating the soma of two adjacent L2/3 excitatory neurons (primate: 19 axons, 40 synapses; mouse: 57 axons, 151 synapses). (**B**) *Inset:* shared (green lines) and unshared axons (black lines) between neighboring neuron pairs. Scatter plot of the number of axons synapsed with the soma and perisoma of neighboring neuron pairs versus the proportion of those axons shared between the two neurons (primate: 0.40 ± 0.02 shared axons, n = 60 axons making 162 synapses across 3 neuron pairs; mouse: 0.21 ± 0.02 shared axons, n = 188 axons making 504 synapses across 3 neuron pairs; P=0.1). (**C**) *Inset*: the number of shared axons between a central neuron (blue/red circle) and all neighboring neurons plotted against their distance from the central soma (*d*). Scatter plot showing two analyses for each species (primate: P1, n = 10 axons, 145 synapses, 36 neurons, P2, n = 9 axons, 103 synapses, 36 neurons; mouse: M1, n = 32 axons, 455 synapses, 49 neurons, M2, n = 25 axons, 397 synapses, 49 neurons). Scale bar = 15 µm. Data: mean ± SEM. *P*-values: two-tailed Mann-Whitney U test.

### Energy Constraints and Scaling Artificial Neural Networks

Finally, we hypothesized that the main factor driving the observed differences in neuronal connectivity between mouse and primate was the metabolic cost of maintaining synapses as the number of neurons increases with brain size. To test this idea, we trained recurrent neural networks (RNNs) with separate excitatory/inhibitory units preforming a cognitive task common to both mouse and macaque visual systems: delayed match-to-sample task (Dudchenko, 2004; Masse et al., 2018; Yang et al., 2019; Zhang et al., 2019) (**Fig. 6A** and see **Methods**). We hypothesized that metabolic costs would provide strong evolutionary pressure (Bullmore and Sporns, 2012) for the sparser connectivity we observed in primate neurons, particularly as the number of neurons in primate brains is estimated to be 90 times more than mouse (∼6.4 billion in *Macaca mulatta*, ∼71 million in *Mus musculus*) (Herculano-Houzel et al., 2015; Herculano-Houzel et al., 2007; Herculano-Houzel et al., 2006). We focused on two forms of ‘cost’: one related to somatic firing rates (*activity cost; AC*) and one on building/maintaining synapses (*synapse cost; SC*). Importantly, while AC was activity-dependent, SC was not (i.e. the proliferation of synapses was independent of their activity or AC (see Methods)).

**Fig. 6.**
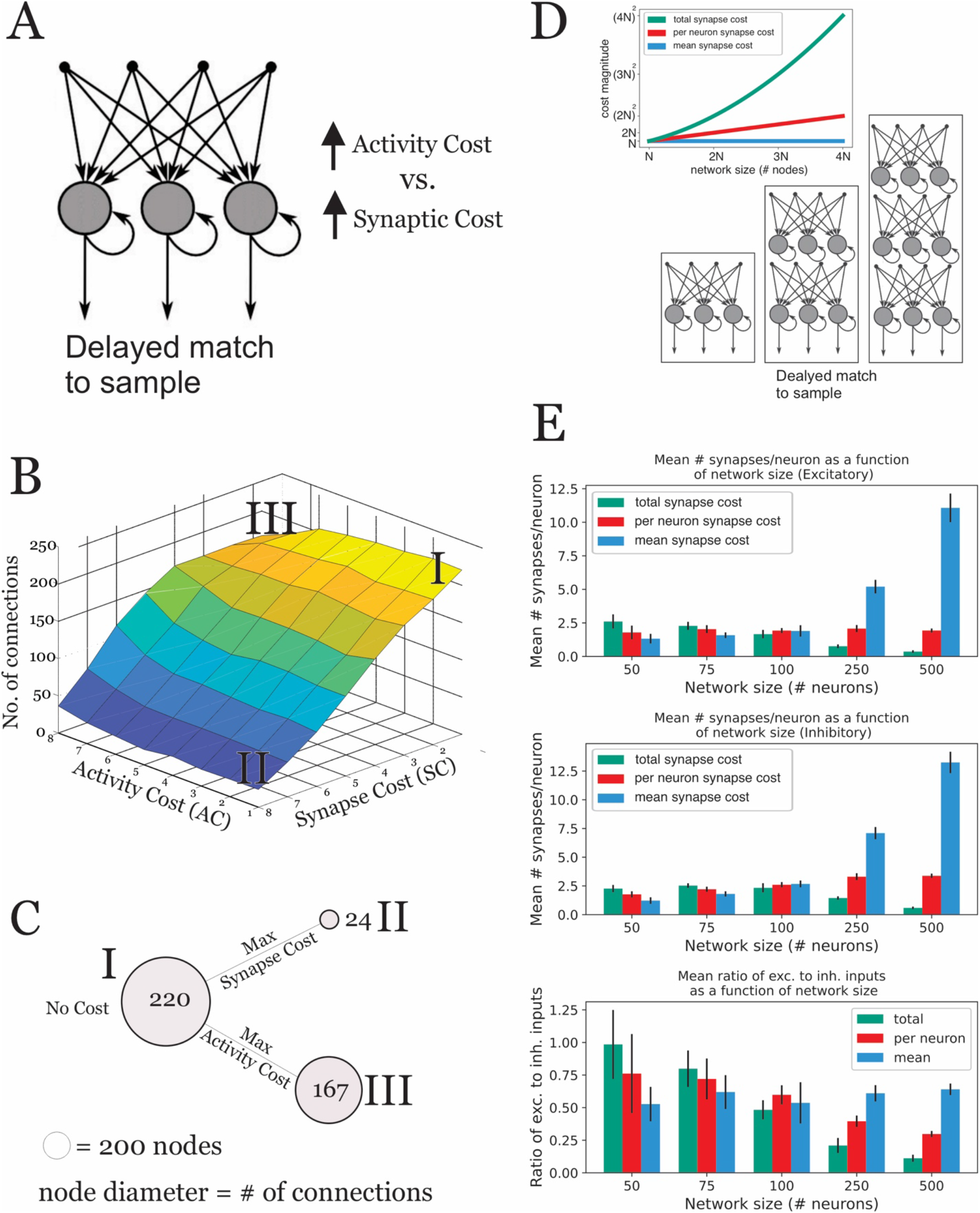
Metabolic costs on firing rates and synaptic machinery regulate connection density in artificial recurrent neural networks. **(A) Schematic of the recurrent neural network.** A matrix of recurrent neural networks (RNNS) consisting of 200 nodes (i.e. neurons) were constructed each with a different activity and synapse cost imposed while trained to perform a delayed match to sample cognitive task. **(B)** 3D heatmap of the connection density of RNNs trained to perform a cognitive task under different costs. For each combination of activity cost weighting (penalty on mean firing rate; AC) and synapse cost weighting (penalty on mean connection strength; SC), 10 networks were trained to perform the same delayed match-to-sample task. Roman numerals mark extreme points in the matrix: (I) no AC and SC cost, (II) max SC, no AC cost, and (III) max AC, no SC cost. **(C)** Cartoon summarizing RNN results when AC or SC are maximally applied alone. Each circle represents the 200 nodes used for the initial cognitive task in (A), and the size of the circle represents the final number of synapses the network has at the end of the training. With no cost (I; SC nor AC) the network has a mean 220 synapses. With a maximum SC cost and no AC cost (II), the network has a mean 24 synapses. With a maximum AC cost and no SC cost (III), the network has a mean 167 synapses. **(D)** *Inset:* line graph of cost magnitude versus network size. Different models of tabulating cost scale differently as network size increases: total synapse cost scales quadratically, synapse cost per neuron scales linearly, and mean synapse cost does not scale. *Cartoon*: RNNs with different numbers of nodes (i.e. neurons) with a fixed activity cost were trained to perform a delayed match to sample cognitive task. **(E)** *Top/middle*: Mean number of synapses per neuron (+/− std. dev.) as a function of network size, shown separately for excitatory neurons (top) and inhibitory neurons (middle). *Bottom*: Mean ratio of excitatory to inhibitory inputs per neuron as a function of network size (+/− std. dev.). In all cases, n=10 networks per comparison.

We performed two sets of experiments: First, we trained RNNs of fixed numbers of nodes with a range of different AC/SC cost combinations (n=10 networks per AC/SC combination, 200 nodes) and measured the average numbers of connections per node at the end of training. We find that while increasing AC and SC reduces the number of connections per node after training, the effects of SC on synapse number were far more pronounced than AC (**Fig. 6B-C**, **S7A**). An index of dispersion (IoD) analysis of the mean synapse count per neuron confirmed this conclusion (see **Methods**). When SC magnitude is fixed and AC magnitude varies, IoDs of mean synapse count are significantly lower than the converse, fixed AC and varied SC (**Fig. S7B**; P=9.39e-28, Mann-Whitney U test) than when SC magnitude varies and AC magnitude is fixed. Further analysis of networks with large AC costs revealed that activity is minimized predominantly through increases in the numbers of inhibitory connections, unlike the sparsity of inhibitory connections found in the non-human primate brain (**Fig. S7C-D**).

We thus focused our second experiment on the effects of SC as the size of networks increases. Specifically, we experimented with 3 models of tabulating the ‘cost’ of synapses in networks of increasing size: (1) total synapse cost – which scales quadratically with the number of nodes, (2) synapse cost per neuron - which scales linearly with the number of neurons, and (3) mean synapse cost, used in experiments described above (Fig. 6A-C) and which does not scale with network size (**Fig. 6D**). As networks increase in node size (from 50 to 500 neurons), total synapse penalty produced networks with reductions in excitatory and inhibitory connections and E/I ratio as seen in the primate EM data, decreasing by a factor of 5.13 as the number of neurons increased by an order of magnitude (i.e. from 50 to 500) (**Fig. 6E**, green bars; P=5.52e-21, Wald test). Networks penalized instead on synapse number per neuron or mean synapse strength either held constant the number of E synapses and increased the number of I synapses or actually increased the numbers of both E and I connections (**Fig. 6E**, red bars, P=6.20e-45; blue bars, P=3.61e-07, Wald test). We conclude that as neural networks scale in size, energy constraints on building and maintaining the total number of connections results in sparse networks as observed in our primate versus mouse EM results.

## Discussion

We leveraged recent advances in automated serial EM to compare the largest volume ‘connectome’ collected to date of non-human primate primary visual cortex with morphologically similar neurons in the mouse primary visual cortex. Our results directly demonstrate, for the first time, that both excitatory and inhibitory primate neurons in L2/3 receive far less spine, and soma synapses than their mouse counterparts (**Table S1**) but retain basic properties of neuronal wiring (i.e. the number of shared axonal somatic inputs on neighboring neurons). We extend previous work limited to measuring bulk synaptic density in neuropil (i.e. synapses/mm^3^) (Ascoli et al., 2008; DeFelipe et al., 2002; DeFelipe et al., 1999; Hsu et al., 2017; Medalla and Luebke, 2015; Peters et al., 2008; Sherwood et al., 2020), by detailing where changes in different types of synapses occur on non-human primate and mouse excitatory and inhibitory neurons (e.g. axons, distal dendrites, etc.; **Fig. 7A**), and how those changes affect how neurons wire together. We report the first analyses classifying and comparing the ratio of excitatory and inhibitory *onto* inhibitory neurons dendrites and somas across species (**Fig. 2A**) as such data is difficult to infer from light microscopy and likely requires EM volumes large enough to accommodate large fractions of neurons. As both primate inhibitory and excitatory neurons maintain sparse synaptic connections relative to mouse counterparts, we conclude the importance of sparse connectivity in primate neuronal functioning. Finally, using neurobiologically-inspired artificial neural networks trained on cognitive tasks with metabolic constraints, we demonstrate that energy costs of maintaining more connections as the numbers of nodes (neurons) increases drives the sparsity of neuronal connections *in silico*.

**Figure 7.**
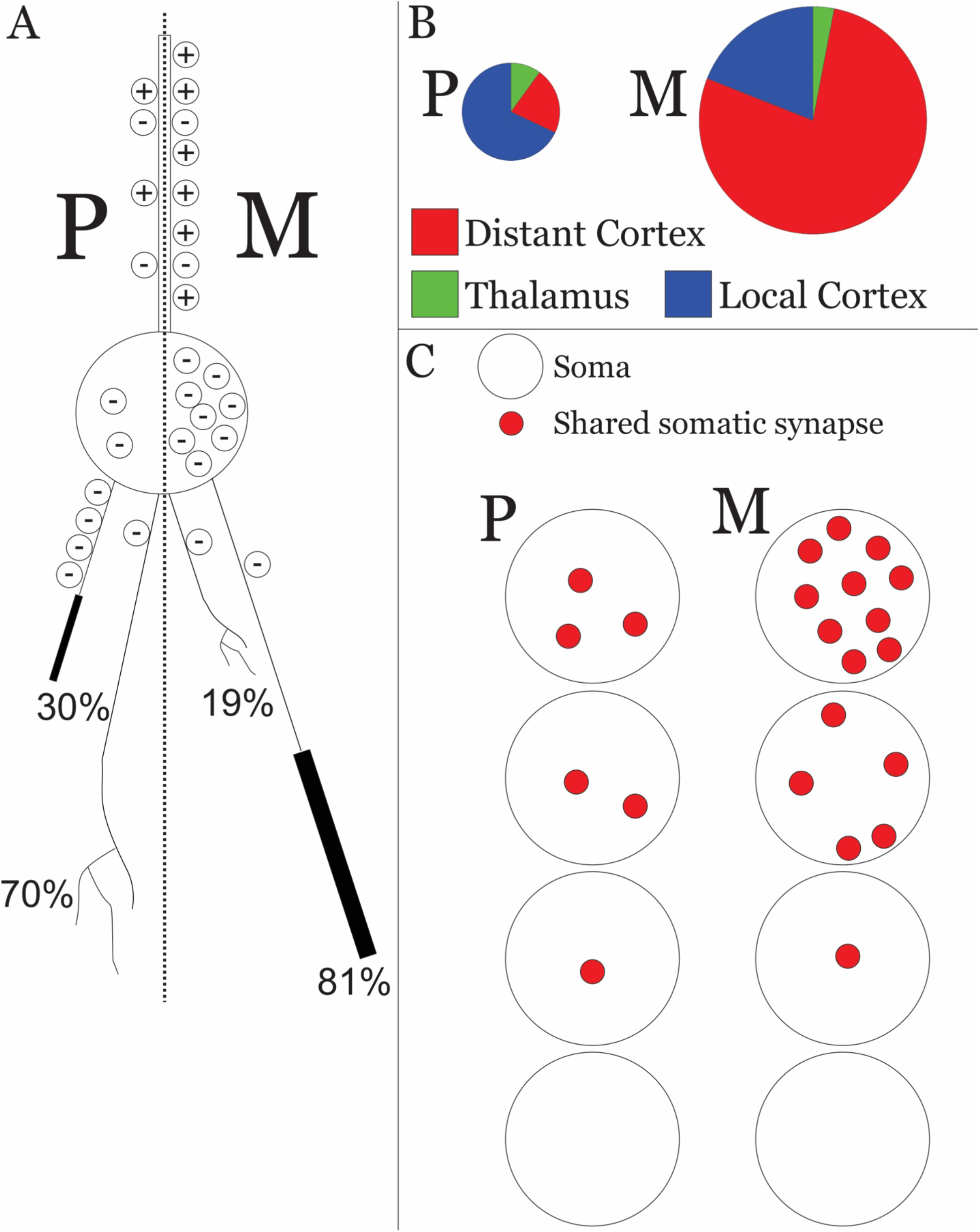
Summary and interpretation of major findings. **(A)** Cartoon depicting prototypical primate (P) and mouse (M) L2/3 excitatory neuron. Along their dendrite, primate excitatory neurons have fewer excitatory inputs (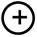) than mouse but an equivalent number of inhibitory inputs (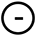) along their shaft. Primate neurons have fewer inhibitory inputs onto their somas (large circle) relative to mouse. Most L2/3 primate excitatory neurons have axons that locally branch (70%) compared to axons that myelinate (black bar) shortly after leaving the soma. (30%), and myelinated axons have more axoaxonic synapses compared to axons that locally branch. Most L2/3 mouse excitatory neurons myelinate (81%) compared to those that locally branch (19%). Locally branched and myelinated axons receive an equivalent number axoaxonic synapses in mouse excitatory neurons. **(B)** Putative proportion of distant, thalamic, and local inputs received by L2/3 excitatory primate (P) and mouse (M) neurons. The diameter of each circle represents the overall total inputs each species neuron receives. Primate excitatory neurons receive proportionally more local inputs whereas mouse receive proportionally more distant connections. Primate neurons also receive more thalamic inputs than mouse neurons. **(C)** Cartoon showing primate excitatory neurons receive fewer total somatic inputs from inhibitory axons than mouse neurons, but primate and mouse neurons have a similar decrease in axonal sharing as a function of distance from a given soma.

There are several potential limitations to the current study. First, although we have limited numbers of animals (1 mouse and 2 non-human primates), the effect sizes recorded were large, statistically significant, and remarkably consistent across neurons and animals. Indeed, given that we saw limited variability across neurons for these effects across species and that some of our findings in mice are consistent with published reports (Gilman et al., 2017; Iascone et al., 2020; Schneider-Mizell et al., 2020; Turner et al., 2020a; Yoshimura and Callaway, 2005), we are confident that these results reflect real differences in the connectivity of primate and mouse neurons in V1, L2/3. Finally, the sparsity of primate connections relative to the mouse might be a feature of primary visual cortex. Indeed, downstream higher order cortical areas, for example prefrontal cortex, have been hypothesized to have a high degree of recurrent connectivity to support the higher cognitive functions localized there (Pollen, 2003). In such areas one might expect to see the opposite trend (i.e. denser connectivity in primate vs. mouse).

The second limitation is our definitions of ‘adult’ mouse and primate. With little literature about developmental staging of neuronal connections across species, we chose standard definitions (i.e. beyond known critical periods and prior to age-related cognitive decline). Indeed, future experiments detailing how neurons connect over both developing, ‘adult’, and aged animals will help establish timelines for comparing connectivity across these and other species. One possible outcome could be that the differences we see in ‘adult’ brains are either differences in initial synaptic densities at birth or different rates of synaptic addition and pruning over development.

### Redistributions of synapses on single neurons across species

We find connectomic evidence for changes in the distributions of types of synapses innervating individual non-human primate and mouse neurons. For excitatory innervation, our data suggests that primate cortical neurons are more strongly influenced by thalamus and local cortical innervation than mouse cortical networks (**Fig. 7B**). First, primate neurons have less spines than mouse counterparts but a larger fraction of those spines contain a second inhibitory input, an indication that the primary excitatory spine input is from thalamus (see **Fig. 3D, S4, and Fig. 7B**) (Kubota et al., 2007; Rodriguez-Moreno et al., 2018). Second, since primate axons myelinate less and more likely form local collaterals than mouse neurons (**Fig. 4A**, bottom panel, **7A**, **Table S2**), and since L2/3 neurons mediates cortical-cortical communication (Sarid et al., 2015; Winguth and Winer, 1986), a simple inference would be that primate cortical neurons are more connected with local cortex and mouse neurons receive more input from distal cortical regions (**Fig. 7B**).

We also find changes in the relative distribution of the types of inhibitory synapses across primate and mouse neurons. Despite receiving substantially fewer numbers of somatic inhibitory inputs and reduced excitatory synapses, both primate excitatory and inhibitory neurons maintain similar numbers of dendritic shaft synapses (**Fig. 2**, **3, Table S1**). Additionally, primate excitatory neurons have increased numbers of second inhibitory synapses on spines (**Fig. 3D, S4, Table S1**). As a result, primate neurons have lower E/I ratios than mouse neurons (i.e. they receive proportionally more inhibitory inputs than excitatory) reflecting the increased influence of inhibitory connections on dendritic trees as opposed to the soma (**Fig. 2A**, **3A-D**, **Fig. 7A**). These redistributions could result from species differences in the prevalence of interneuron types in the primate brain or the presence of novel inhibitory cell types (Džaja et al., 2014; Hodge et al., 2019; Krienen et al., 2020; Sultan and Shi, 2018). An emphasis on dendritic inhibition in non-human primates might support species specific dendritic processing in primate neurons as recently reported (Gidon et al., 2020).

### The Sparsity of Inhibitory Connections Across Species

In contrast to changes in dendritic inhibition, the innervation patterns of somatic targeting inhibitory axons were remarkably similar across species despite mouse excitatory neurons receiving many more somatic inhibitory synapses. Across species, interneuronal connectivity was neither dense nor non-specific (Fino et al., 2013; Packer and Yuste, 2011) but instead sparse: neurons only tens of microns apart were unlikely to ‘share’ the same pre-synaptic inhibitory axon (**Fig. 5C**, **7C**). Indeed, we found many instances where inhibitory axons that synapse on some soma, come in direct physical contact with other neuronal soma and dendrites without innervating these neurons in either species (data not shown). These results instead favor models where interneurons connect with precision and selectivity (Yoshimura and Callaway, 2005). One consequence of non-human primate neurons receiving sparse excitatory innervation and dominated by inhibition is that in such networks, it is easier for any internally generated variability to be uncorrelated between neurons (Stringer et al., 2016).

### Functional Implications

Finally, detailed maps of mouse and primate V1 neuronal connections could help explain similarities and differences in the response properties of mouse and non-human primate neurons. For instance, neurons in both species perform edge detection and feature extraction, (Huberman and Niell, 2011; Van Hooser, 2007) but mouse V1 neurons often have larger receptive field sizes, ‘coarser’ tuning curves, and reach maximal firing rates much lower (∼2x) than primate V1 neurons (Wang et al., 2016) for both visual stimuli and optogenetic manipulations (Sanzeni and Histed, 2020; Sweeney and Clopath, 2020). A simple potential explanation from our data suggests that feature extraction could be mediated by the pattern of inhibitory innervation, relatively similar across species, but that the magnitude of responses are encoded by the absolute number of connections, substantially different across species. Previous simulations/models comparing mouse and primate neurons have concluded that even morphologically similar neurons with different electrical properties could account for these different physiological responses (Segev, 1992). The sparsity of connections on primate neurons will likely inform and extend these models (Achard and Bullmore, 2007; Eyal et al., 2014; Papo et al., 2014; van den Heuvel et al., 2016) for explaining functional differences in response properties of neurons across species.

### Energy Constraints and Scaling Natural and in silico Neural Networks

For in silico computations, while there has seen increasing success in solving problems such as image recognition and language understanding using artificial neural networks, scaling these networks to billions of parameters (i.e. synapses) has benefited from the idea of “conditional computation” in which a given stimulus only activates a sparse set of a (large) model, thereby making it both possible and beneficial to train such extremely large models while requiring similar amounts of data and compute (Shazeer et al, 2017). While the idea that energy budgets drive potential evolutionary changes in brains is not new (Attwell and Laughlin, 2001; Harris et al., 2012), we suggest a specific hypothesis for future experimentation: that the number of connections in brains decreases as the number of neurons increase as consequence of the energy burden of maintaining synapses. For example, this work predicts that in the visual systems of animals intermediate in neuronal number between primate and mouse, (e.g. the ferret, rat, or cat) the number of E and I connections on individual neurons would also be intermediate to the measurements listed here. Our work suggests as brains scale in size and number of neurons, neurons reduce and redistribute their connectivity profiles (**Fig. 7A-B**), changing some network properties like local versus distal excitatory connections and conserving others like the sparsity of somatic innervation. (**Fig. 7C**).

## Acknowledgments

We thank J.H.R. Maunsell, S.M. Sherman, V. Jain, and V. Sampathkumar for discussions; J.H.R. Maunsell and V. Jain for valuable comments on the manuscript; Marek Niekrasz and entire ARC team for assistance with animal care; H. Li and K Norwood for computer infrastructure and coding; Tom Uram at Argonne National Laboratory for assistance with implementing alignment code on Argonne’s supercomputer.

## Author contributions

N.K. and G.A.W. conceived of the experiment and performed analyses; G.A.W. collected the data (sample preparation, imaging, image alignment, and analyses; N.K. and G.A.W. wrote the manuscript. D.P. plotted Fig. 1D and S2B. M.R. Rosen performed all *in silico* RNN experiments (Fig. 6, S7, S8).

## Data and materials availability

The raw EM data and Matlab code used for data analysis is freely available upon request.

## Methods

### Animals

A wild-type (C57BL/6) male mouse 15 weeks old and two wild type male Rhesus Macaque, ages 11 years old and 14.5 years old were used in this study. Animal care and perfusion procedures were followed according to animal regulations at the University of Chicago’s Animal Resources Center (ARC) and approved IACUC protocols.

### Data Acquisition

Unless otherwise noted, brains for primate and mouse were prepared in the same manner and as previously described (Hua et al., 2015). Briefly, an anesthetized animal was first transcardially perfused with 0.1 M Sodium Cacodylate (cacodylate) buffer, pH 7.4 (Electron microscopy sciences (EMS) followed by fixative containing 2% paraformaldehyde (EMS), 2.5% glutaraldehyde (EMS) in 0.1 M Sodium Cacodylate (cacodylate) buffer, pH 7.4 (EMS). Volumes were adjusted for each animal: for mouse, we used 10 ml of cacodylate buffer followed by 20 ml of fixative; for primate, we used 1 L of cacodylate buffer followed by 6 L of fixative. The brain was removed and placed in fixative for at least 24 hours at 4°C. For the primate brain, a small posterior region of the brain was cut away using a surgical scalpel and V1 was identified using anatomical landmarks (*e.g.* calcarine sulcus) and comparing with a published atlas (Paxinos, 2009). A 300 µm vibratome section encompassing V1 was removed and put into fixative for 24 hours at 4°C. For mouse, a 300 µm vibratome section encompassing V1 was removed and put into fixative for 24 hours at 4°C. Mouse V1 was identified using the Allen Brain Institute reference atlas (http://mouse.brain-map.org/experiment/thumbnails/100048576?image_type=atlas). Brain slices were washed extensively in cacodylate buffer at room temperature and stained sequentially with 2% osmium tetroxide (EMS) in cacodylate buffer, 2.5% potassium ferrocyanide (Sigma-Aldrich), thiocarbohydrazide, unbuffered 2% osmium tetroxide, 1% uranyl acetate, and 0.66% Aspartic acid buffered Lead (II) Nitrate with extensive rinses between each step with the exception of potassium ferrocyanide. The sections were then dehydrated in ethanol and propylene oxide and infiltrated with 812 Epon resin (EMS, Mixture: 49% Embed 812, 28% DDSA, 21% NMA, and 2.0% DMP 30). The resin-infiltrated tissue was cured at 60°C for 3 days. Using a commercial ultramicrotome (Powertome, RMC), the cured block was trimmed to a 0.8 x 1.5 mm or 0.8 x 2.4 mm rectangle for mouse and primate brain samples, respectively. 3,000, 40nm thick sections were collected from each block on polyamide tape (Kapton) using an automated tape collecting device (ATUM, RMC) and assembled on silicon wafers as previously described (Kasthuri et al., 2015). The serial sections were acquired using backscattered electron detection with a Gemini 300 scanning electron microscope (Carl Zeiss), equipped with ATLAS software for automated wafer imaging. For 40nm and 6nm resolution data sets, sections were brightness/contrast normalized and rigidly aligned using TrakEM2 (FIJI) followed by non-linear affine alignment using AlignTK on Argonne National Laboratory’s super computer, Cooley. Different image processing tools were packaged into Python scripts that can be found here: https://github.com/Hanyu-Li/klab_utils.

### Data Analysis

Aligned datasets were manually skeletonized and annotated using the publicly available software, Knossos (https://knossos.app). Neurons were reconstructed by first identifying the soma followed by tracing their dendritic arbors. Classes of cell types were identified by distinguishing anatomical properties: Excitatory neurons by the presences of dendritic spines, interneurons by the lack of spines and numerous shaft synapses, glial cells by their extensive branching, lack of spines and synapses, and cytoplasmic granules. Skeleton information was exported into tab delimited matrices using homemade Python scripts that compute skeleton features from the Knossos xml annotation file. Code is freely available here: https://github.com/knorwood0/MNRVA. Quantification and plotting of different anatomical features were performed in Matlab and excel. Two-tailed Mann Whitney U statistics test was used to test for significance (Marx et al., 2016) between mouse and the aggregate of primate 1 and primate 2 datasets. Mouse, primate 1, and primate 2 datasets were individually plotted to visualize the similarity between primate samples. Below is a brief description of each quantification reported:

Figure 1:

1. Inter-soma distance: The position of every soma in the L2/3 high resolution (6nm) dataset was marked in Knossos as a node sized to the full 3D volume of the soma, and every pairwise Euclidian distance between the center of mass of every soma was calculated.
2. Branches per neuron: All dendrites were fully reconstructed in every neuron in the L2/3 high resolution (6nm) dataset (see Supplementary Figure 2). Dendritic length and number were extracted from skeletons using automated scripts (see MNRVA code in above link).

Figure 2, 3, S4:

1. Spine and shaft synapse frequency: the dendrites of neurons were divided into an average of 15, ∼10µm segments. The number of spine or shaft synapses were counted manually within each segment and divided by the actual segment length to calculate spine synapses/µm and shaft synapses/µm. Synapses were identified by the presence of a post-synaptic density and vesicles on pre-synaptic axon. The distance between the furthest point of the 10µm segment and the soma was also computed using the Euclidian distance formula to calculate the distance from the soma.
2. E/I ratio for excitatory neurons: we used the following equation: E/I = total spine synapses / (total shaft + total soma/perisoma + total second spine synapses). Total values for excitatory and inhibitory inputs were derived by calculating:

Excitatory: total spine = spine synapse density * average total dendritic length
Inhibitory: total shaft = shaft synapse density * average total dendritic length; total soma/perisoma = total soma + perisoma synapse number; total 2^nd^ spine synapse = 2^nd^ spines/spine * total spine
3. E/I ratio for inhibitory neurons (Fig 3D): we divided the number of excitatory shaft synapses by inhibitory synapses. Axons that primarily (> 75%) made synapses on shafts of excitatory or inhibitory neurons, soma, and 2^nd^ spines were classified as inhibitory axons. Axons that primarily (> 75%) made synapses on primary spine heads or excitatory neurons and the shafts of inhibitory neurons were classified as excitatory axons.

Figure 4:

1. 4A: Excitatory neurons were classified by tracing their axons and scored for whether they myelinated (MA) or branched (uMA). Neurons with axons that left the dataset before they myelinated or branched were not scored.
2. 4C: the axon initial segment (AIS) length was calculated as the Euclidian distance from the soma to the furthest axoaxonic synapses from the soma.
3. 4D-E: Axoaxonic synapses were identified as symmetric synapses between axonal boutons containing vesicles and a post-synaptic density positioned on the axon being analyzed.

Figure 5:

1. 4B: All somatic synapses were first identified and marked in Knossos between two pairs of neighboring excitatory neurons that were approximately in the center of the 6nm dataset and as physically close to each other as possible (∼10 µm from center of soma mass). These somatic synapses were used as seed points for fully reconstructing inhibitory axons and every synapse these axons made with both pairs of neurons was annotated for the type of synapse (soma, perisoma, shaft, 2^nd^ spine). The proportion of axons that synapsed with both neighboring neurons was then calculated from the axon reconstructions to determine how many axons were shared between the two neurons.
2. 4C: In Knossos, all somatic synapses were identified and marked on a soma that was approximately in the center of the 6nm dataset. These somatic synapses were used as seed points for fully reconstructing inhibitory axons and every synapse these axons made was annotated for the type of synapse (soma, perisoma, shaft, second spine) and which neuron in the FOV it synapsed with. The number of axons that connected with the central soma and another neuron, and the inter-soma distance between the central soma and neuron that shared an axon was calculated. Inter-soma distance was also calculated between the central soma and neurons that received zero synapses from these inhibitory axons.

Supplementary Figure 5:

1. S5A: The length of inhibitory axons was calculated as described above (3C) and the number of branches were manually counted in Knossos
2. S5B: Skeleton nodes in Knossos were placed over the bouton and adjusted to approximate the bouton size. The node radius was used to calculate the surface area using S.A = π4r^2^
3. S5C: The length of inhibitory axons used in Figure 4 was computed using homemade scripts (https://github.com/knorwood0/MNRVA) and the number of synapses were determined from Knossos manual annotations
4. S5D: The number of axons that innervated the soma was manually counted from reconstructions used in Figure 4.
5. S5E: the number of soma synapses from inhibitory axons was derived from reconstructions and synapse annotations used for Figure 4.

### Artificial recurrent neural networks

Rate-based artificial recurrent neural networks (RNNs) were trained to perform a delayed match-to-sample (DMS) task. This task, developed in primate neurophysiology experiments, requires the comparison of sequentially-presented dot motion stimuli to indicate a directional match. Networks consisted of excitatory and inhibitory units obeying Dale’s law (4 excitatory units for each inhibitory unit). Synapses in these networks were also subject to short-term synaptic plasticity according to the method previously described (Masse et al., 2018), with 50% of synapses briefly facilitated by presynaptic activity and 50% briefly depressed. Training occurred through supervised learning; network parameters (weights and biases) were adjusted using the backpropagation-through-time algorithm to minimize the categorical cross-entropy loss *L(ŷ*, *y*) between predicted output *ŷ* and true output *y*. Weights were initialized randomly using a gamma distribution and constrained to be strictly positive; as a synapse’s weight is pushed below 0 in magnitude during training, it is effectively pruned, ceasing to affect network output and receiving no further adjustment. All networks were trained to stable, comparable levels of performance (>90% accuracy on the DMS task) (Fig S8). After training, synapses were counted as the number of outgoing weights with positive magnitude. Networks were trained in Python using the Tensorflow framework; code implementing this training process, along with all relevant hyperparameters for reproduction, is available on Github.

### Metabolic costs

Metabolic costs on neural activity and synapse size were implemented through components added to the cross-entropy loss function used during RNN training. The activity cost (AC) is given by 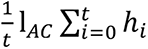, the mean of all neurons’ activity across all timepoints of a trial. The synapse cost (SC) is given by 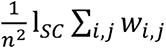, the mean synapse weight between all pairs of a network’s *n* neurons. Two other variants of SC were also used (Fig. 6D-E): 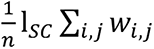, a penalty on the total synapse weight per neuron in a network, and I*_SC_* Σ*_i,j_ w_i,j_*, a penalty on the total weight of all synapses in the network. Each of these costs was multiplied by a constant factor, λ, titrating that cost’s relative strength during training. Increasing λ_AC_ increased the strength of the activity cost during training, constraining networks more strongly to adopt solutions with lower average activity; similarly, increasing λ_SC_ constrained networks more strongly to adopt solutions with lower synaptic weights. The maximal values of λ*_AC_* and λ*_SC_* tested and reported here were chosen based on their compatibility with stable network training. After increasing λ*_AC_* and λ*_SC_* no longer yielded networks that could consistently be trained past 90% accuracy on the DMS task (Fig. S6), corresponding with inappropriately strong regularization, λ*_AC_* and λ*_SC_* were capped, with subsequent analyses restricted to groups of networks with λ*_AC_* and λ*_SC_* below the threshold of instability.

---

**Fig. S1.**
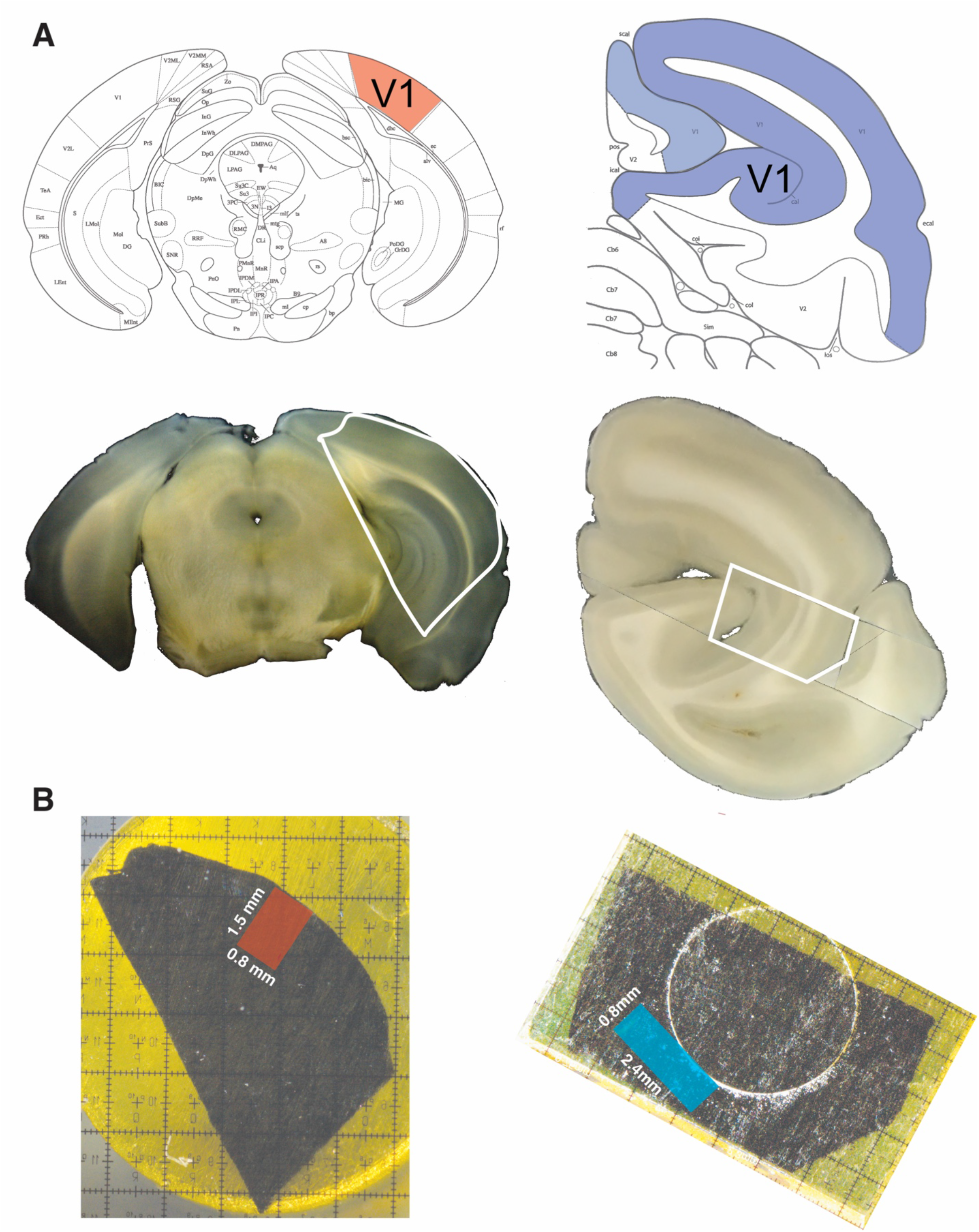
EM tissue processing of mouse and primate V1. (**A**) For mouse (left), a whole brain was vibratome sectioned into 300 µm thick coronal sections and a section containing V1 was identified using anatomical landmarks from a reference atlas. For primate (right), a small posterior piece of brain was removed with a scalpel and then vibratome sectioned into 300 µm thick coronal sections and a section containing V1 was identified using anatomical landmarks from a reference atlas. For both species, a smaller piece that spanned all cortical layers of V1 was cut from the coronal section with a scalpel (white outline). (**B**) The smaller section containing V1 was prepared for EM (see **Methods**) and further trimmed to a final region containing all cortical layers of V1 (0.8 mm x 1.5 mm, red box in mouse, 0.8 mm x 2.4 mm, blue box for primate) for serial sectioning.

**Fig. S2.**
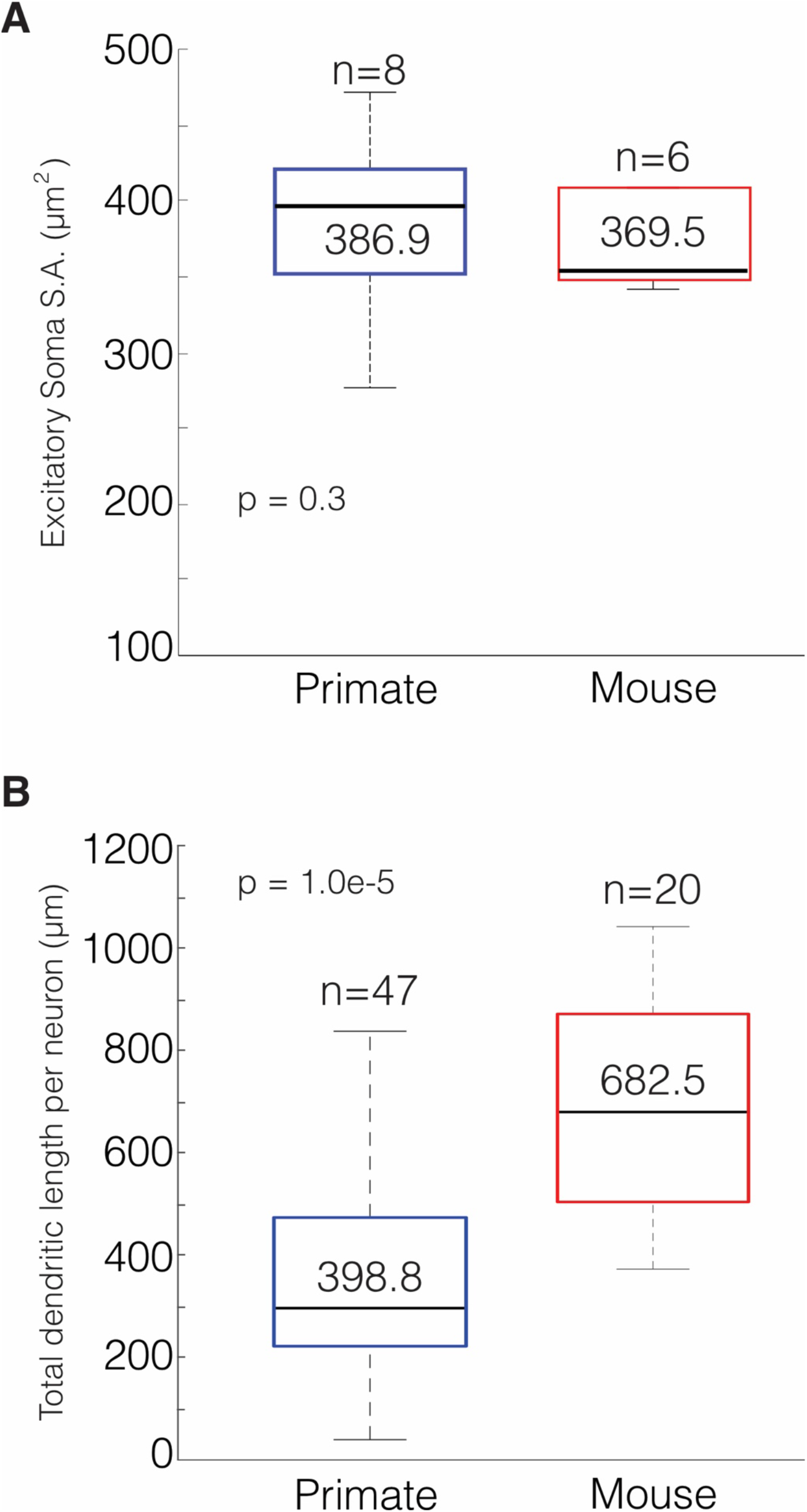
Primate and Mouse L2/3 excitatory neuron soma surface area and total dendritic branch length. (**A**) Box-and-whisker plot of soma surface area (S.A.) of primate and mouse neurons (primate: 386.9 ± 20.8 um^2^, n = 8 neurons; mouse: 369.5 ± 12.4 um^2^, n = 6 neurons; P = 0.3). (**B**) Box-and-whisker plot of total dendritic length per neuron (primate: 398.8 ± 41.2 µm, n = 47 neurons; mouse: 682.5 ± 45.9 µm, n = 20 neurons; P = 1.0e-5). Data: mean ± SEM. *P*-values: two-tailed Mann-Whitney U test.

**Fig. S3.**
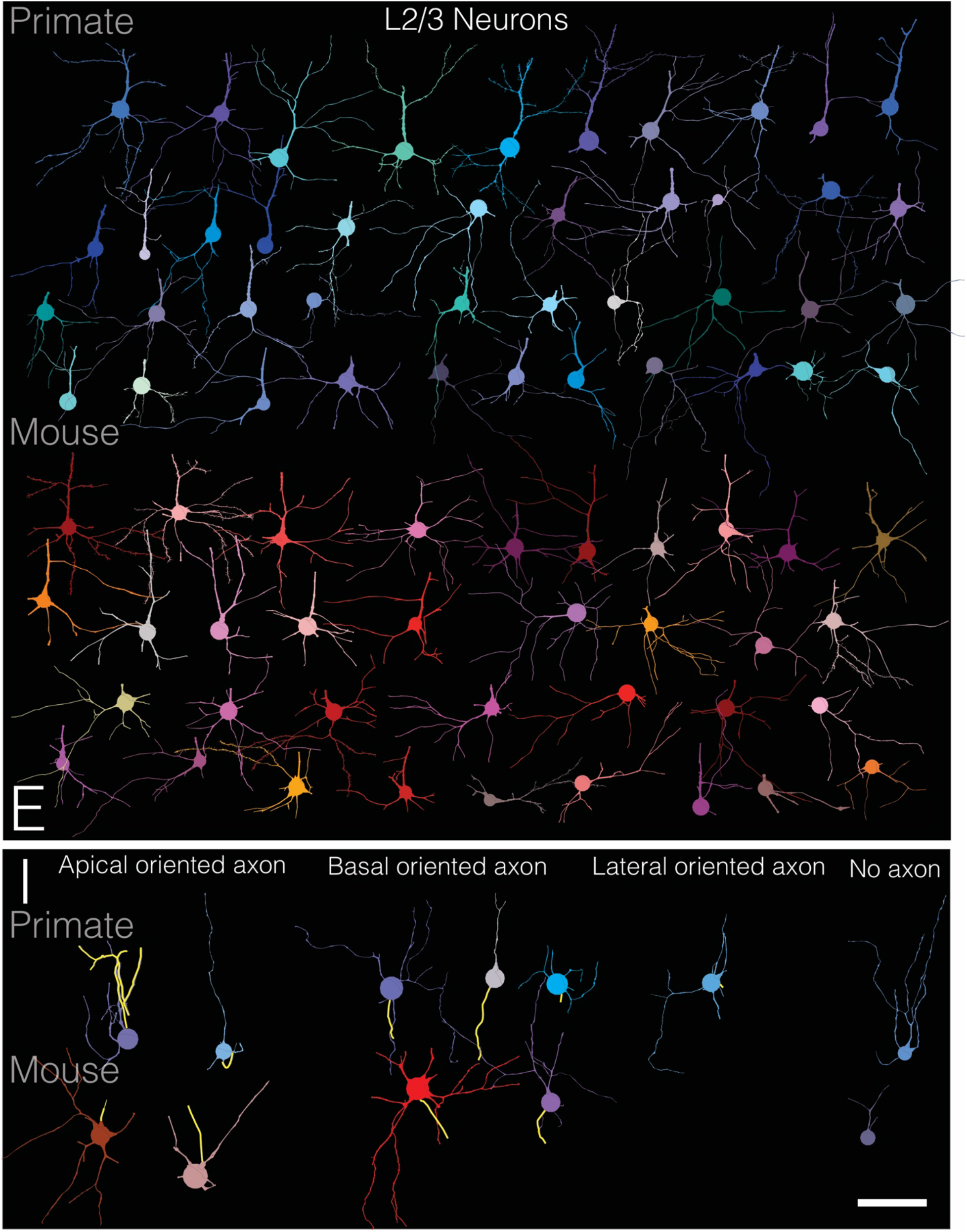
Rogues’ gallery of V1, L2/3 mouse and primate neurons reconstructions. Fine resolution (∼ 6nm XY resolution) 3D reconstructions of excitatory (E) and inhibitory (I) neurons from primate and mouse L2/3 used for population level analyses. 69 primate and 42 mouse neurons had their dendritic arbors fully reconstructed. Top panel, excitatory (E) neurons for each species. Bottom panel, inhibitory (I) neurons were subdivided by the orientation of their axon (highlighted in yellow). Reconstructions lacking an axon were at the edge of the dataset and thus missing the axon-containing portion of the neuron. Scale bar = 50 µm.

**Figure S4.**
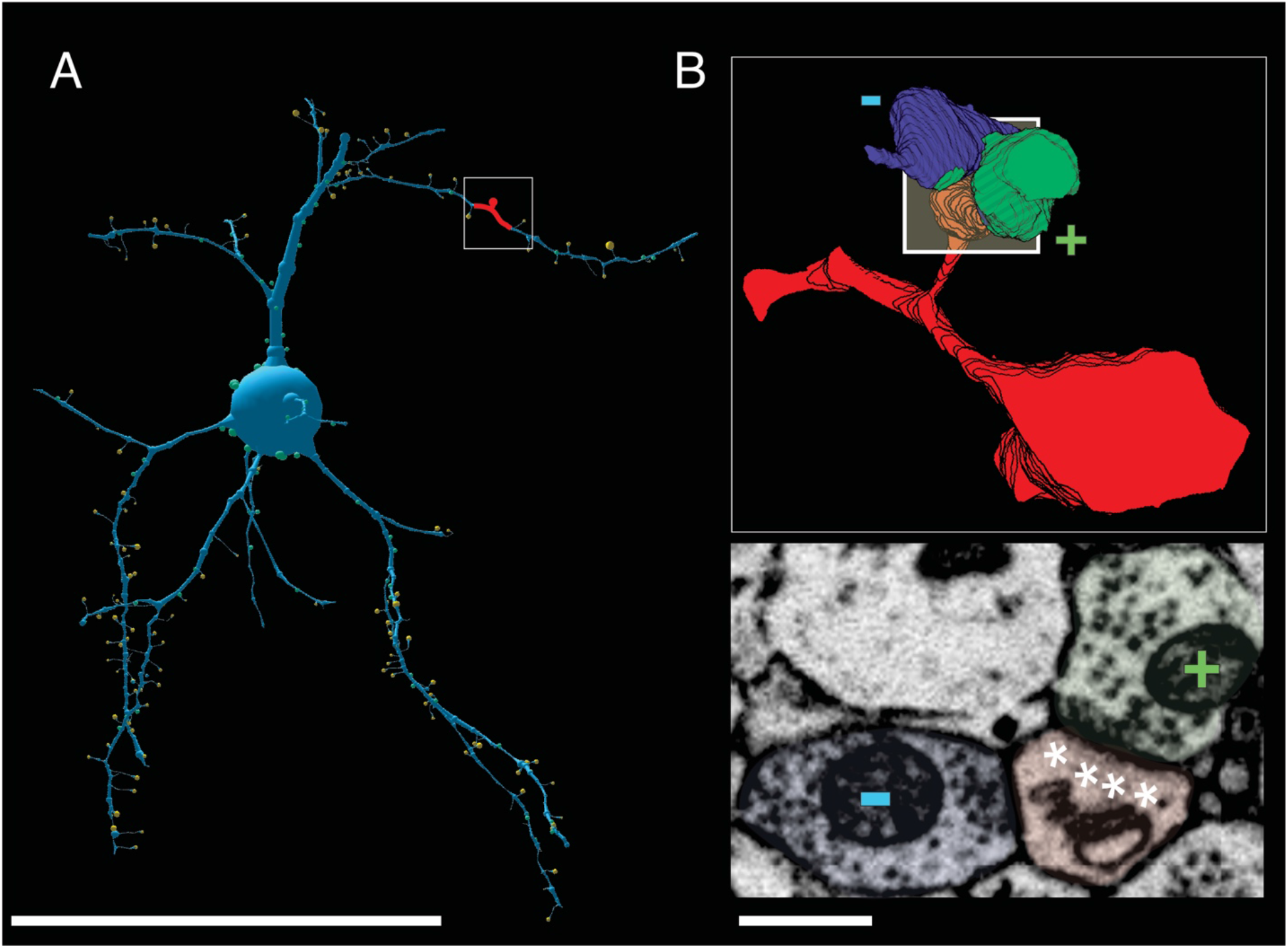
Primate excitatory neurons have more 2^nd^ spine synapse. **(A)** Reconstruction of a representative primate excitatory neuron. White box with red spine depicts location of representative spine with a second synapse. **(B)** 3D reconstruction (*top*) and matched 2D EM (*bottom*) image of a primate dendritic spine (red) with a 2^nd^ synapse: an excitatory synapse on the spine head (green/+) and inhibitory synapse on the side (blue/-). Asterisks in middle panel show the post-synaptic density on the (+) synapse. Scale bar = 50 µm (A), 500 nm (B).

**Figure S5.**
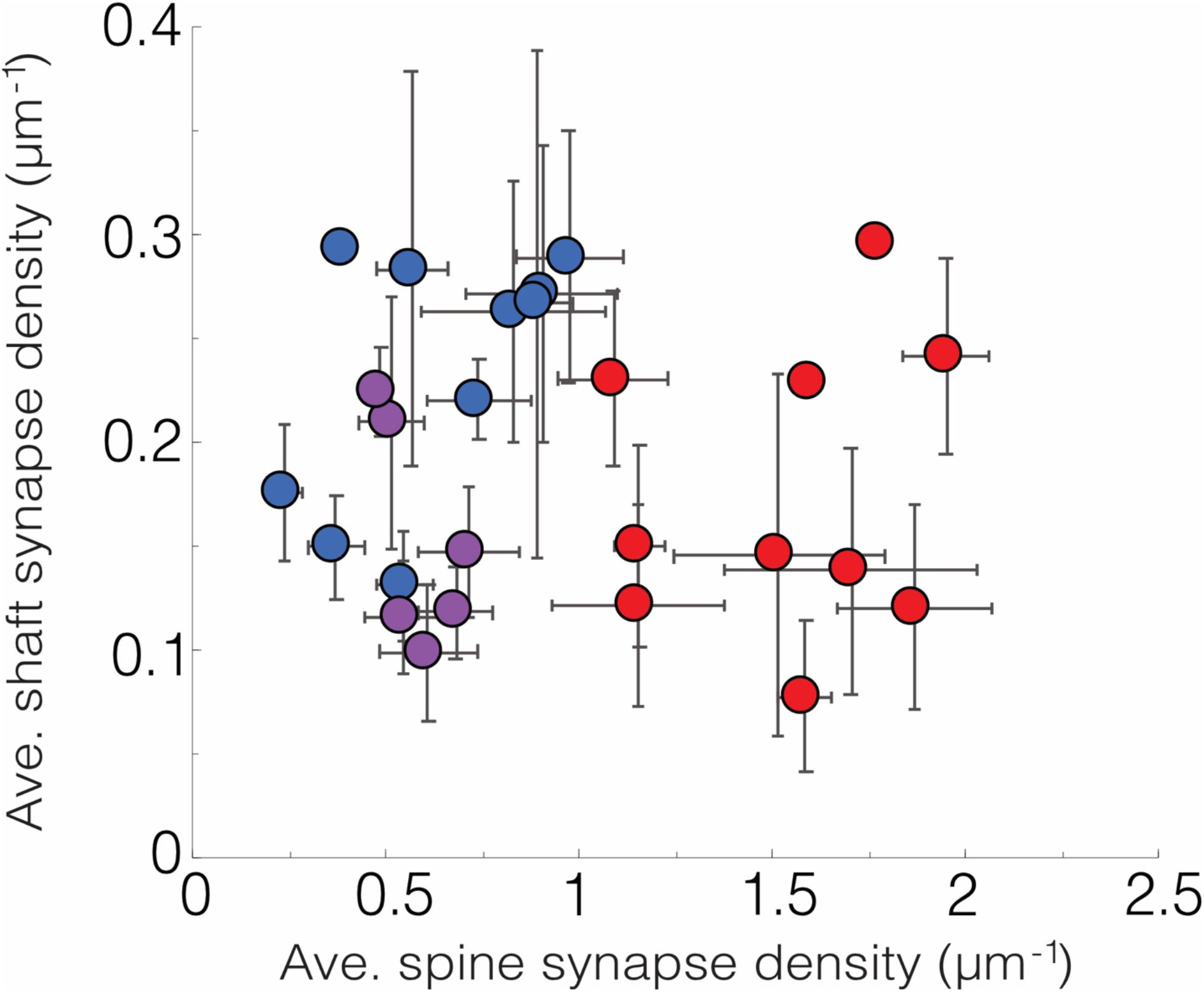
Excitatory neuron variability is lower within species and between species. Scatter plot of the average number of shaft synapses/µm versus average number of spine synapses/µm of individual mouse and primate neurons (primate 1- primate 2 variance = 0.04; primate 1 (or 2) – mouse variance = 0.28).

**Fig. S6.**
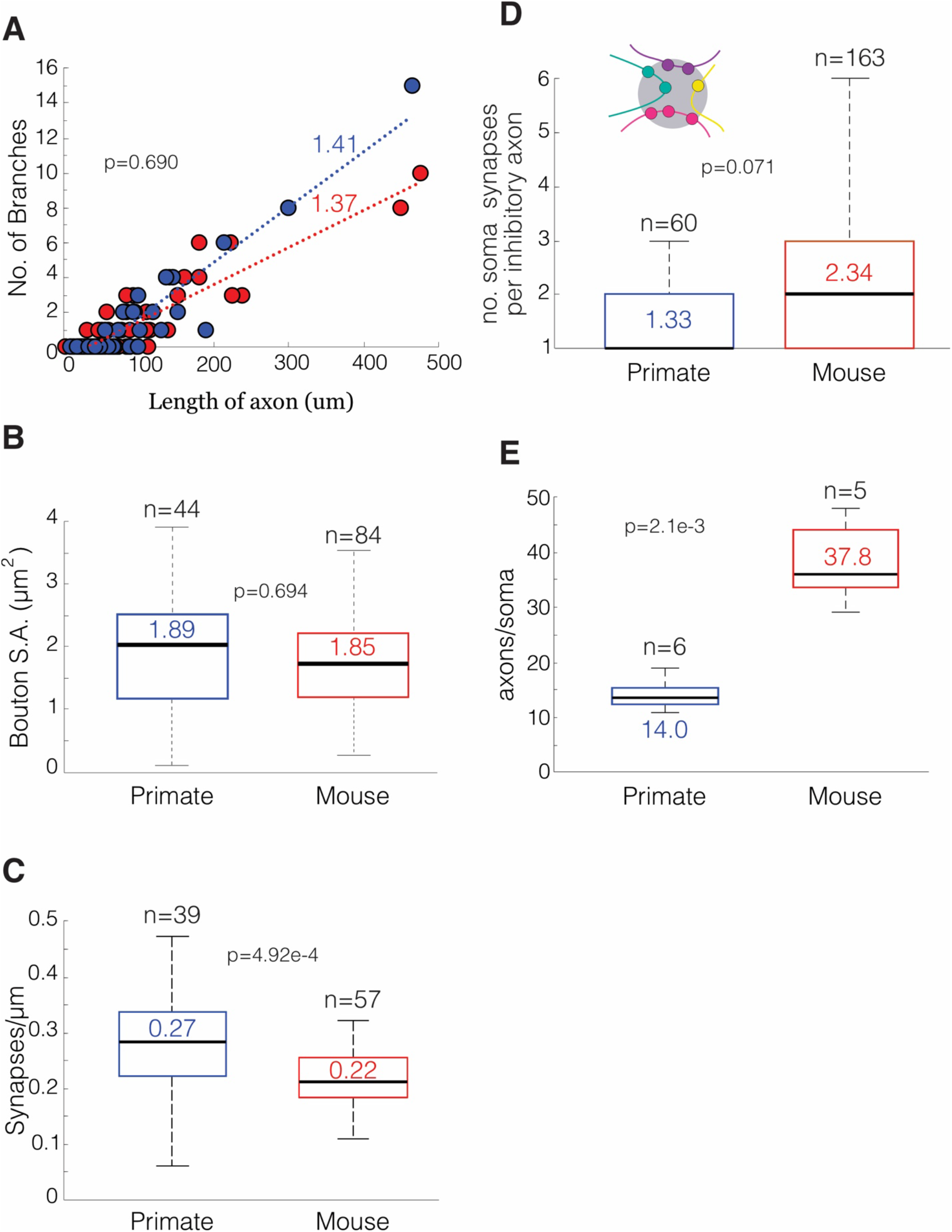
Individual anatomical properties of inhibitory axons are unchanged between primate and mouse. (**A**) Scatter plot of the number of branches versus length of inhibitory axons (mean primate: 1.41± 0.46 branches/µm, n = 39 axons; mean mouse: 1.37 ± 0.29 branches/µm, n = 57 axons; P=0.690). (**B**) Box-and-whisker plot of the bouton surface area of inhibitory axons at the sight of somatic synapses on excitatory neurons in primate and mouse (primate: 1.89 ± 0.14 um^2^, n = 44 boutons across 2 neurons; mouse: 1.85 ± 0.10 um^2^, n = 84 boutons across 2 neurons; P = 0.694). (**C**) Box-and-whisker plot of the number of synapses/µm an inhibitory axon makes in primate and mouse (primate: 0.27 ± 0.02 synapses/µm, n = 857 synapses across 39 axons; mouse: 0.22 ± 0.01 synapses/µm, n = 1118 synapses across 57 axons; P = 4.92e-4). (**D**) Box-and-whisker plot of the number of soma synapses per individual inhibitory axon on primate and mouse neurons. *Inset*: cartoon showing individual axons depicted as different colored lines making synapses (circles) on the soma of a neuron (primate: 1.33 ± 0.15 soma synapses/axon, n = 115 synapses across 60 axons; mouse: 2.34 ± 0.11 soma synapses/axon, n = 381 synapses across 163 axons; P = 0.071). (**E**) Box-and-whisker plot of the number of axons innervating the soma in primate and mouse (primate: 14.0 ± 1.0 (axons/soma, n = 84 axons across 6 soma; mouse: 37.8 ± 2.5 axons/soma, n = 198 axons across 5 soma, P=2.1e-3). Data: mean ± SEM. *P*-values: two-tailed Mann-Whitney U test.

**Figure S7.**
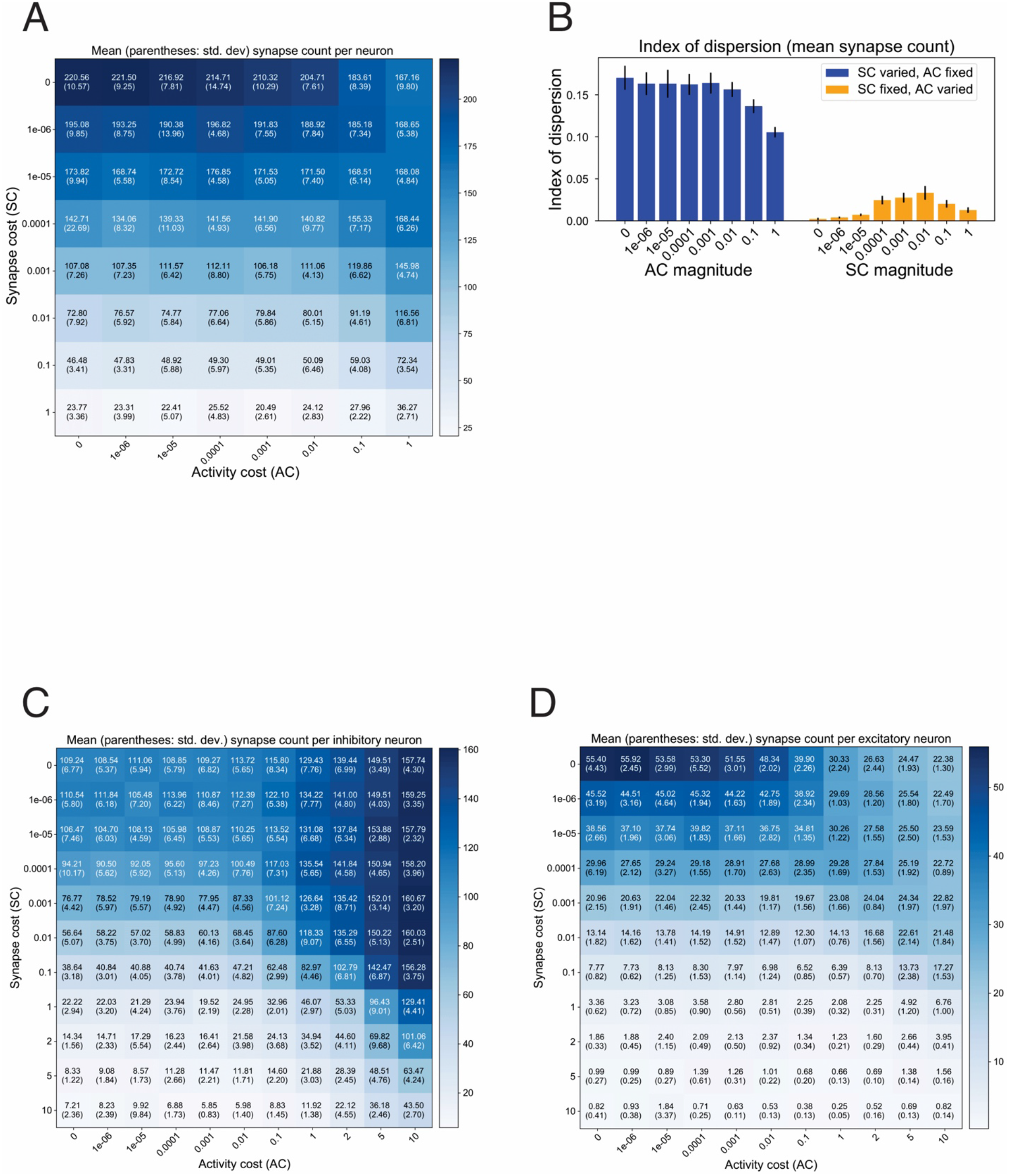
Metabolic costs on firing rates and synaptic machinery regulate connection density in artificial recurrent neural networks. **(A)** Heatmap of same results from Figure 6A but showing numerical outputs of RNNs performance of the cognitive task. Each cell in the heatmap reports the mean number of synapses per neuron in these networks, +/− standard deviation. (**B)** Indices of dispersion (windowless Fano Factor) of mean synapse count per neuron. Blue bars (left) show column-wise indices of dispersion, when AC is fixed but SC is varied (n=80 networks per group; mean=0.15, std. dev.=0.02); orange bars (right) show row-wise indices of dispersion, when AC is varied but SC is fixed (n=80 networks per group; mean=0.016, std. dev.=0.012). **(C)** Mean (+/− std. dev.) synapse count per excitatory neuron across a range of activity/synapse cost combinations. Average taken over n=10 networks, each with 200 neurons. **(D)** Mean (+/− std. dev.) synapse count per inhibitory neuron in the same set of networks shown in B.

**Figure S8.**
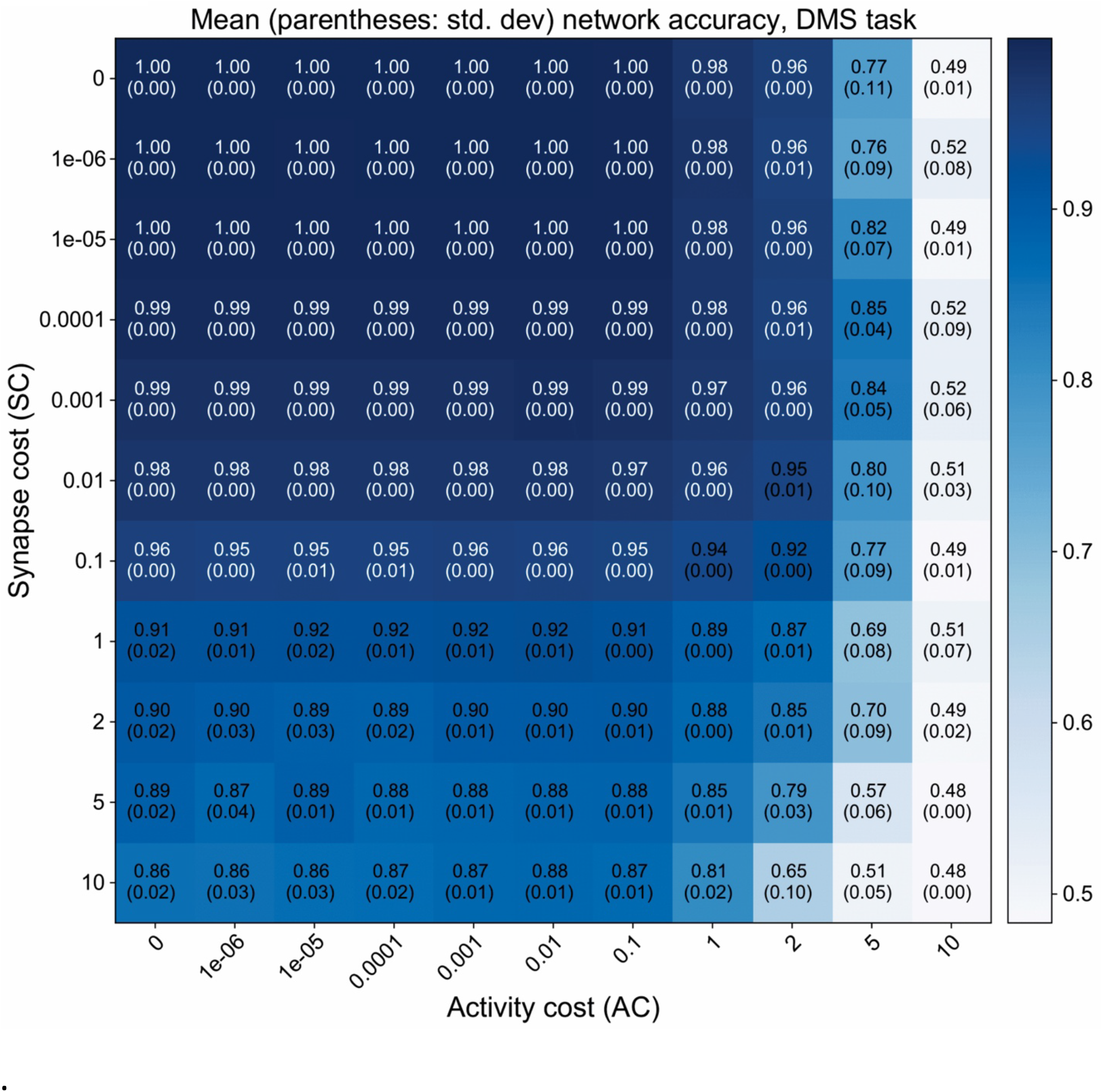
Mean (+/− std. dev.) of network accuracy across all nets trained to perform the DMS task, across a variety of weightings of activity and synapse costs. n=10 networks were trained per cost combination, each with 200 nodes.

**Table S1.** Summary table listing quantifications from this report. When noted with an asterisk, the anatomical feature was quantified from one mouse and one primate dataset; otherwise data was quantified from one mouse and 2 primate datasets. Statistical significance was calculated using the Mann-Whitney test. n.c. = no change between mouse and primate datasets.

|  | Mouse<br>(n = 1 brain) | Primate<br>(*n = 1 brain)<br>(n = 2 brains) | n<br>ms =mouse<br>pr =primate | p-value | Fold change<br>in primate |
| --- | --- | --- | --- | --- | --- |
| **Measurement (mean ± SEM)** |  |  |  |  |  |
|  | **Excitatory Neurons** |  |  |  |  |
| *Soma density ($\mu\text{m}$) | 53.3 ± 0.4 | 47.6 ± 0.3 | ms: 62 neurons<br>pr: 71 neurons | 2.8e-3 | n.c. |
| *Soma surface area ($\mu\text{m}^2$) | 370 ± 12.4 | 387 ± 20.8 | ms: 6 neurons<br>pr: 8 neurons | 0.3 | n.c. |
| *Branch number/neuron | 18.0 ± 0.9 | 16.2 ± 1.2 | ms: 20 neurons<br>pr: 47 neurons | 0.08 | n.c. |
| *total dendritic length ($\mu\text{m}$) | 683 ± 46 | 399 ± 41 | ms: 20 neurons<br>pr: 47 neurons | 1e-5 | 1.7x lower |
| Spine synapse/$\mu\text{m}$ | 1.34 ± 0.1 | 0.56 ± 0.03 | ms: 82 10$\mu\text{m}$ dendrite fragments on 10 neurons<br>pr: 174 10$\mu\text{m}$ dendrite fragments on 16 neurons | 1.5e-14 | 2.4x lower |
| Shaft synapse/$\mu\text{m}$ | 0.18 ± 0.02 | 0.18 ± 0.01 | ms: 82 10$\mu\text{m}$ dendrite fragments on 10 neurons<br>pr: 16 10 $\mu\text{m}$ dendrite fragments on 16 neurons | 0.23 | n.c. |
| Synapse/(peri)soma | 102 ± 5.3 | 25 ± 1.9 | ms: 388 synapses on 6 neurons<br>pr: 237 synapses on 13 neurons | 7.4e-5 | 4.1x lower |
| 2nd Spine synapses (%) | 3.0 ± 0.0 | 13.0 ± 0.0 | ms: 180 spines on 3 neurons<br>pr: 437 spines on 7 neurons | 6.6e-4 | 4.3x **higher** |
| Est. total excitatory synapses | 5846 ± 626 | 1790 ± 281 | ms: 770 spines on 4 neurons<br>pr: 669 spines on 7 neurons | 6.0e-3 | 3.3x lower |
| Est. total inhibitory synapses | 1048 ± 22 | 759 ± 59 | ms: 109 shaft, 401 soma synapses on 4 neurons<br>pr: 207 shaft, 173 soma synapses on 7 neurons | 6.0e-3 | 1.4x lower |
| E/I | 5.55 ± 0.5 | 2.42 ± 0.4 | (combined total of Est. total excitatory and Est. total inhibitory synapses) | 6.1e-3 | 2.3x lower |
|  | **Inhibitory Neurons** |  |  |  |  |
| Shaft synapse/$\mu\text{m}$ | 2.7 ± 0.1 | 1.7 ± 0.1 | ms: 39 10$\mu\text{m}$ dendrite fragments on 4 neurons<br>pr: 58 10$\mu\text{m}$ dendrite fragments on 10 neurons | 3.31e-8 | 1.6x lower |
| Synapse/(peri)soma | 248 ± 15 | 44 ± 7 | ms: 743 synapses on 3 neurons<br>pr: 307 synapse on 7 neurons | 0.012 | 5.6x lower |
| E/I | 9.6 | 2.6 | ms: 32 axons traced off of 1 neuron<br>pr: 54 axons traced off of 2 neurons | n.d. | 3.7x lower |
|  | **Somatic innervating axons** |  |  |  |  |
| *Branches/$\mu\text{m}$ | 1.4 ± 0.3 | 1.4 ± 0.5 | ms: 57 axons<br>pr: 39 axons | 0.7 | n.c. |
| *Bouton surface area ($\mu\text{m}$) | 1.9 ± 0.1 | 1.9 ± 0.1 | ms: 84 boutons on 2 neurons<br>pr: 44 boutons on 2 neurons | 0.7 | n.c. |
| *Synapses/$\mu\text{m}$ | 0.22 ± 0.0 | 0.27 ± 0.0 | ms: 1118 synapses on 57 axons<br>pr: 857 synapses on 39 axons | 4.9e-4 | n.c. |
| *Peri(soma) synapses/axon | 2.3 ± 0.1 | 1.3 ± 0.2 | ms: 381 synapses on 163 axons<br>pr: 115 synapses on 60 axons | 0.07 | n.c. |
| *Axons/soma | 38 ± 3 | 14 ± 1 | ms: 198 axons on 5 soma<br>pr: 84 axons on 6 soma | 2.1e-3 | 2.7x lower |
| *% shared (nearest neighbor) | 21 ± 0.0 | 40 ± 0.0 | ms: 188 axons, 504 synapses on 3 neuron pairs<br>pr: 60 axons, 162 synapses on 3 neuron pairs | 0.1 | n.c. |

n.c. = no change

**Table S2.** Summary table of axon initial segment synapse differences. Summary of results from quantifications of mouse and primate L2/3 excitatory neurons with unmyelinated, locally branching axon (uMA) and those that myelinate ∼25µm from the soma and do not branch (MA) in the imaged field of view. Statistical significance was calculated using the Mann-Whitney test. n.a. = not applicable and n.c. = no change between mouse and primate datasets.

| **Mouse**<br>(n=1 brain) | MA | uMA | n | p-value | Fold change |
| --- | --- | --- | --- | --- | --- |
| Estimate % population | 81% | 19% | MA: 13 neurons<br>uMA: 3 neurons | n.a. | 4x fold more<br>MA neurons |
| AIS length ($\mu\text{m}$) | 23 ± 1 | 28 ± 2 | MA: 13 neurons<br>uMA: 3 neurons | 0.4 | n.c. |
| # AIS synapses | 9 ± 1 | 14 ± 2 | MA: 121 synapses on 13 neurons<br>uMA: 41 synapses on 3 neurons | 0.08 | n.c. |

| **Primate**<br>(n=2 brains) | MA | uMA | n | p-value | Fold change |
| --- | --- | --- | --- | --- | --- |
| Estimate % population | 33% | 67% | MA: 13 neurons<br>uMA: 3 neurons | n.a. | 2x fold more<br>uMA neurons |
| AIS length ($\mu\text{m}$) | 22 ± 1 | 19 ± 1 | MA: 13 neurons<br>uMA: 3 neurons | 0.13 | n.c. |
| # AIS synapses | 16 ± 2 | 4 ± 0.6 | MA: 121 synapses on 13 neurons<br>uMA: 41 synapses on 3 neurons | 1.54e-7 | 4x fold more synapses<br>on MA neurons |

Measurement (mean ± SEM)
n.a. = not applicable n.c. = no change

